# Sleep forms flexible context representations in toddlers

**DOI:** 10.64898/2026.06.16.732120

**Authors:** Lisa Bastian, Eva-Maria Kurz, Lilli Gutjahr, Hannes Noack, Jan Born

**Affiliations:** Institute for Medical Psychology and Behavioral Neurobiology, University of Tübingen, Germany; Max Planck School of Cognition, Leipzig, Germany; Department of Child and Adolescent Psychiatry, Psychosomatics and Psychotherapy, University Hospital of Psychiatry and Psychotherapy, University of Tübingen, Germany; German Center for Diabetes Research (DZD), Tübingen, Germany; Center for Integrative Neuroscience, Tübingen, Germany; German Center for Mental Health (DZPG), Tübingen, Germany; Institute for Diabetes Research and Metabolic Diseases of Helmholtz Center Munich at University of Tübingen (IDM), Tübingen, Germany

**Keywords:** Context memory, Sleep, Memory consolidation, Spindle-slow oscillation coupling

## Abstract

Sleep consolidates episodic memory through a hippocampus-dependent process in adults. Whether and how sleep supports memory consolidation during early life, when hippocampal function is immature, remains unclear. Here, we examined effects of sleep on the consolidation of spatial context, a core component of episodic memory, in toddlers aged 2 –3 years. Toddlers were familiarized with two spatial contexts, followed by a ∼90-min nap or an equivalent wake period. Afterwards, with a hide-and-seek game we tested their ability to relocate toys within these contexts. Only after post-familiarization sleep, the toddlers showed significant context memory and formed stronger associations between toys and specific contexts compared to wakefulness. Contextual memory was positively correlated with spindle density and slow oscillation–spindle phase-amplitude coupling during non-rapid eye movement (NonREM) sleep. Despite hippocampal immaturity, the sleeping toddler’s brain seems to engage consolidation processes similar to those in adults to form spatial context memory for the flexible use in novel situations.

## Introduction

Sleep supports memory formation through a hippocampus-dependent consolidation process^1–5^. This consolidation process is thought to originate from the repeated replay of newly encoded hippocampal representations during non-rapid eye movement (NonREM) sleep, driven by ripples in the hippocampus and cortex^6^ (∼100 Hz). These ripples are thought to group cofiring within widespread cortical sites during time windows of neural plasticity that are defined by cortical down-to-upstate transitions (i.e. slow oscillations (SOs) with < 1Hz) and local 12–16 Hz spindles, originating from the thalamus^7–10^. The co-occurrence of ripples and spindles, particularly when nested into the excitable SO up-state, are crucial for the long-term storage of memories within neocortical networks^11–14^.

There is a growing body of evidence demonstrating that sleep supports memory formation during early life (e.g.,^15–21)^. However, compared to adults, findings regarding the effects of post-learning sleep versus wakefulness in young children have presented a more mixed picture^22^, which has often been attributed to the immaturity of the hippocampus. Indeed, structural and functional hippocampal development follows a prolonged trajectory^23^, involving processes such as myelination of neuronal axons and ongoing neurogenesis, which continue into later childhood or even adulthood^24–27^. Specifically, subregions of the hippocampus involved in the tri-synaptic pathway suggested to be crucial for forming an episodic memory engram^23^, namely CA3 and dentate gyrus, develop well into the second year at least in rodents^28,29^. Recent volumetric measurements in humans corroborate these findings showing growth of CA3 and dentate gyrus until early adulthood^30^.

Hippocampal immaturity may hamper an optimal functioning also during sleep-dependent consolidation processes^31–33^. Successful memory consolidation relies on effective hippocampal-cortical communication pathways^9^, including central output regions in the entorhinal cortex that develop rather late^34^. Additionally, ripple-spindle coupling as well as SO-spindle coupling has been found to be reduced in juvenile rats and young children, respectively, suggesting that hippocampal-neocortical information transfer underlying memory consolidation during NonREM sleep may not be fully effective^35–40^.

The hippocampus is not only critical for memory consolidation during sleep, but a core structure of the episodic memory system, mainly encoding the spatial context of experienced episodes. Accordingly, the ability to form contextual episodic and spatial memories develops in close conjunction with the maturation of hippocampal functioning^24,41,42^. Although simple short-term associations between events and locations can be formed as early as five months of age^43^, robust spatial context representations emerge only between 1.5 and 2 years^44–48^, and this may depend on sleep. Although behavioral evidence suggests an inflection point for episodic memory formation in the second year of life^49^, these memories are still fragile and easily disrupted well into preschool age^47,50^. We therefore focused on toddlers aged 2–3 years, an age at which spatial context representations are present but not yet adult-like, to ask whether sleep strengthens the consolidation and abstraction of such representations during this early phase of hippocampal development.

Critical to the sleep-dependent consolidation process is that it promotes a gradual transformation of episodic memories into more abstract, generalized representations that reside in neocortical networks and can be flexibly used to integrate novel information in similar situations^3,51^. These abstract neocortical representations can also include spatial context information. In this way, sleep supports the formation of generalized spatial contextual knowledge, which serves as a reference for spatial orientation in similar contexts^52^ (see also^53,54^). In juvenile rats, familiarization with a specific spatial context during infancy enhanced their capability to acutely form object-location memories in the same context when tested as adults^55^. The contextual memory formed during infancy crucially relied on medial prefrontal cortex regions, and at adulthood, was apparently used to enhance spatial behavior, including the formation of novel memory representations.

Here, we used a spatial paradigm adapted from Newcombe et al.^46^ to examine the effects of spindle-SO dynamics during sleep on spatial memory capabilities in toddlers aged between 25 and 34 months, i.e., an age when the hippocampus still shows clear signs of immaturity with spatial and episodic memory functions only starting to emerge. Importantly, the present study was not designed to disentangle the relative contributions of hippocampal versus cortical regions to memory consolidation. While the task and behavioral outcomes are widely considered to rely on hippocampal-dependent context representations, and while sleep spindle and slow-oscillation dynamics are established markers of systems-level consolidation, our design does not allow for direct inferences about the regional origin of the observed effects. Rather, our aim was to examine whether sleep at this developmental stage supports the formation of flexible context representations at the behavioral level and whether this process is associated with canonical sleep oscillations known to mediate hippocampo-neocortical communication.

We first hypothesized that, compared to an equivalent period of wakefulness, a post-familiarization nap would selectively improve contextual aspects of memory, expressed as fewer context errors and a higher Contextualized Memory Index (CMI), without necessarily changing overall accuracy or random errors. Second, based on prior work linking sleep spindles to memory consolidation in children^19^, we expected that higher spindle density during NonREM sleep would be associated with a reduced context error rate and a lower context error rate and a higher CMI, indicating more robust binding of object information to its spatial context. Third, we hypothesized that stronger coupling between slow oscillations and spindles would further predict fewer context errors and increases in CMI, consistent with the view that precisely timed SO–spindle coordination supports the consolidation of abstracted context representations in early childhood.

We find that sleep after familiarization to two spatial contexts of the memory task, in comparison with post-familiarization wakefulness, increased the toddler’s ability to form item-context associations in a subsequent hide-and-seek game, consistent with the idea that sleep helps form abstracted context representations that are not anymore bound to the original familiarization experience. In the hide-and-seek game, the toddlers seemed to use the representations as reference frames for instantly forming novel item-context associations that regulate seeking behavior.

## Results

Each of 28 toddlers (age: 29.46 ± 2.57 months; range 25-34 months, 12 female) was examined in a sleep and a wake condition. In each condition the toddlers were first familiarized with two spatial contexts (i.e. rooms, for photos see **Fig. S3**), each equipped with the same 4 different containers (**Fig. 1a,b**). Familiarization was followed by a 90-min period filled with a nap or wakefulness. Then, the toddlers performed a spatial memory task, i.e., a hide-and-seek game requiring them to hide a puppet in a different container in each room, and after a 5-min delay, to recover the puppets in both rooms (**Fig. 1b**). Memory performance at retrieval was assessed by the numbers of (i) correct trials, i.e., the choice of the correct container in the correct context, (ii) context errors, referring to a correct choice of a container in the wrong context, and (iii) random errors, referring to the choice of one of the 2 containers not hiding a puppet in either context (**Fig. 1c**). Values were transformed into percentages of the number of completed trials (100 %), as for some children not all 4 trials were included in the analysis (see Methods for details).

**Figure 1.**
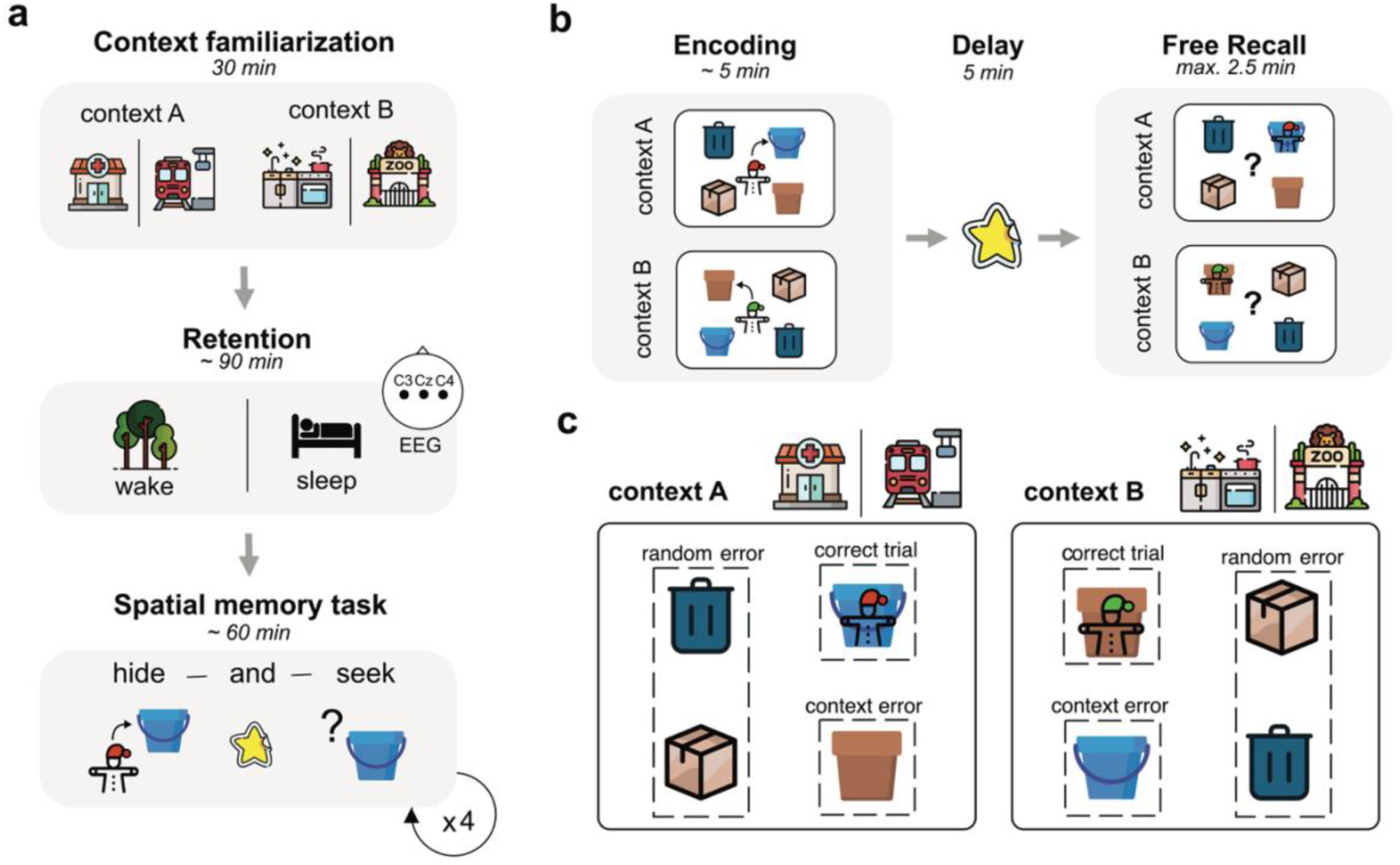
Experimental design, task and behavioral outcome measures. **a**, Each of 28 toddlers was tested on two conditions (sleep vs. wake) at two different locations in the city (lab A vs. lab B) following a within-subject cross-over design. Each condition started with a 30-min context familiarization period in which the participants were familiarized with 2 different rooms each decorated with toys according to a different theme (train station vs. zoo in one condition and teddy clinic vs. play kitchen in the other, see **Fig. S3** for photos). Context familiarization was followed by a ∼90 min retention period filled with a nap or wakefulness. Then, performance on a spatial memory task was tested. **b**, Memory task exemplified for lab B. In each room, 4 different containers were placed. The containers were the same across rooms but arranged differently in each room. The task comprised 4 trials of a hide-and-seek game. In each trial, the child hid a different puppet in one of the four containers, first in one room and then in the other, with the target container differing between the rooms. After a short 5-min delay, the toddler went back into the rooms to look for the puppets and to open the respective container. The room order at encoding and retrieval was counterbalanced across trials (same vs. different context sequence). **c**, Retrieval performance was assessed (across all trials) by the number of correct trials (opening the container with the puppet), context errors (correct container choice but wrong context), and random errors (choice of one of the two containers not hiding a puppet in any of the rooms).

In light of marked performance changes that were observed on this task during early development^46^, we first tested whether our toddlers were able to successfully perform the task. For both, sleep and wake conditions, the percentage of correct trials was above chance level (sleep: t_27_ = 5.984, *P* < 0.001, *d* = 1.13, 95% CI [0.65 – 1.60]; wake: t_27_ = 4.418, *P* < 0.001, *d* = 0.83, 95% CI [0.40 – 1.26]), while random errors were below chance level (sleep: t_27_ = −3.808, *P* < 0.001, *d* = −0.72, 95% CI [−1.13 – −0.30]; wake: t_27_ = −5.531, *P* < 0.001, *d* = −1.00, 95% CI [−1.44 – −0.54]). Further control analyses confirmed no influence of sex or motivation on task performance and a comparable number of completed trials and durations spent in each context in both conditions (see Supplementary Information on “Control analyses” for details).

### Sleep after familiarization reduces context errors

Our main hypothesis stated that sleep following context familiarization facilitates the formation of a spatial context representation. If so, children would commit fewer context errors because they would be better able to distinguish between the two contexts. Interestingly, context error rate was only below chance level for the sleep (*t*_27_ = −1.947, *P* = 0.031, *d* = −0.35, 95% CI [−0.73, 0.04]) but not the wake condition (*t*_27_ = 0.555, *P* = 0.708, *d* = 0.10, 95% CI [−0.2, 0.48]; **Fig. 2b**). We then compared the percentages of correct trials, context errors, and random errors directly between conditions. Linear-mixed effects model analyses (including context sequence and participant age as covariates) confirmed that children committed fewer context errors after sleep compared to the wake condition (**Fig. 2b**; *β =* 0.38, SE *=* 0.18, *P* = 0.033, *d* = 0.41, 95% CI [0.03, 0.73]), indicating that in the sleep condition they less often allocated a correct target container to the wrong room. The benefits of sleep for context separation were particularly important for younger children as they diminished with increasing age (*β* = −0.11, SE *=* 0.05, *P* = 0.034, 95% CI [−0.21, −0.01] for Sleep/Wake x Age interaction, **Fig. 2d**). Sleep did not affect percentages of correct trials (*β = −*0.10, SE *=* 0.17, *P* = 0.570, 95% CI [−0.44, 0.25], **Fig. 2a**) or random errors (*β = −*0.18, SE *=* 0.18, *P* = 0.330, 95% CI [−0.53, 0.18], **Fig. 2c**). Analyses including correct trials and context errors as outcome variables in one model (**Table S5**) confirmed a differential effect of sleep, selectively reducing context errors (*β =* −8.27, SE *=* 3.91, *P* = 0.036, *d* = 0.24, 95% CI [−16.00 – −0.56]) in the absence of an effect on the percentage of correct trials (*β =* 2.72, SE *=* 5.48, *P* = 0.621, *d* = 0.06, 95% CI [−8.10 – 13.54]).

**Figure 2.**
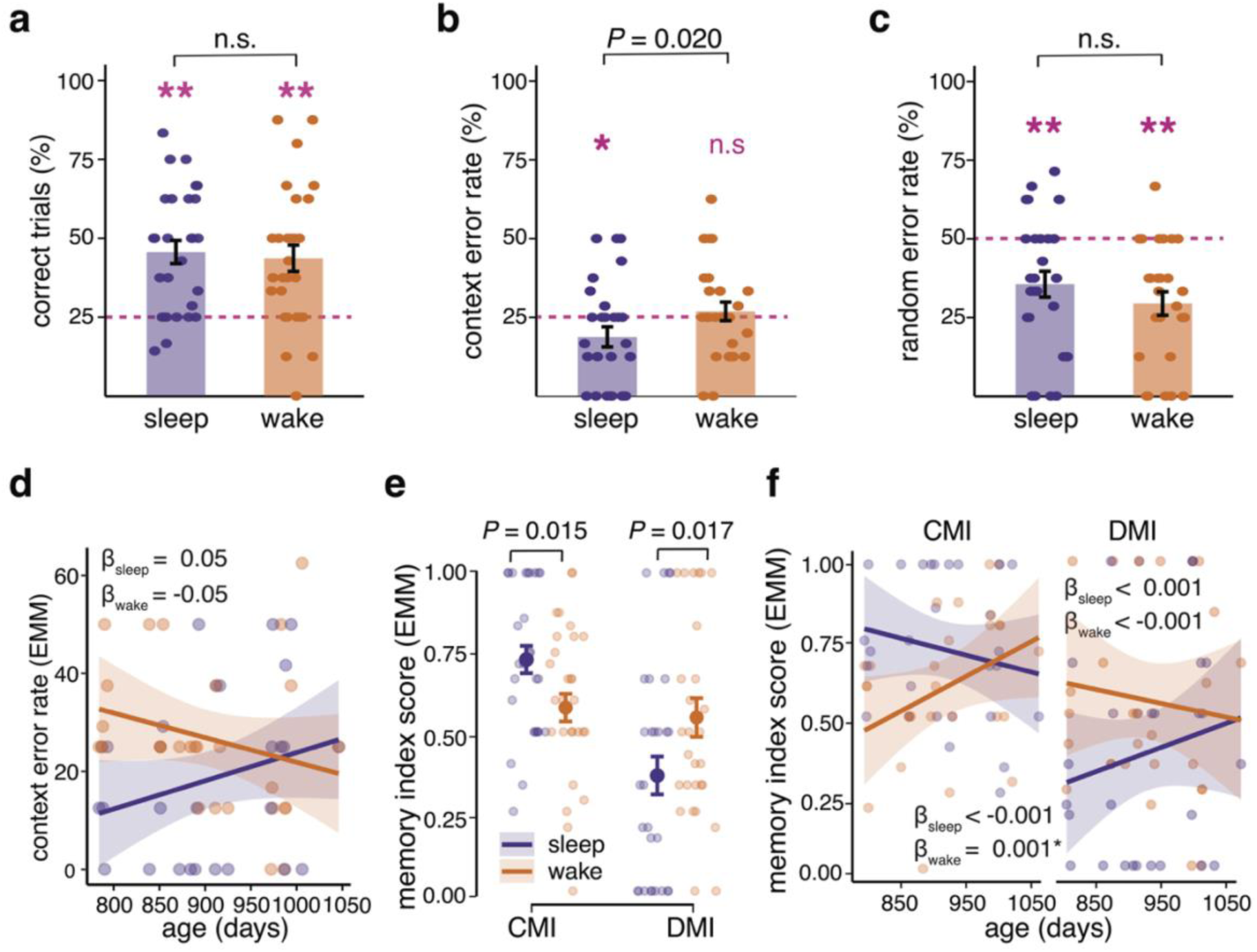
Differences in memory contextualization after sleep and wakefulness. **a**, Percentages of correct trials, **b.** context errors, and **c**. random errors in the sleep and wake conditions (dot plots overlaid, significance for the Condition effect is indicated, N = 28). Asterisks indicate significant differences from chance level (dashed horizontal lines). **d,** Modulation effect of Age on context errors, indicated by estimated marginal means (EMMs) of the Age x Condition interaction for the sleep (purple) and wake (orange) conditions. With increasing age, percentage of context errors increased in the sleep condition but decreased in the wake condition. Slopes are expressed by β coefficients. **e**, Effects on contextualized memory index (CMI) and decontextualized memory index (DMI) indicated as EMMs of CMI and DMI for the sleep (purple) and wake (orange) conditions (dot plots with raw values overlaid, significance for follow-up tests is indicated, N = 27). The CMI is higher than the DMI after sleep but lower than the DMI after wakefulness. **f,** Same as **d** but for the Age x Condition interaction effect on CMI and DMI. \**P* < 0.05, \*\**P* < 0.01.

In addition, this analysis revealed a Sequence effect such that the children made fewer context errors when they entered the two rooms in a different (rather than in the same) sequence during encoding and retrieval (*β =* 11.6, SE *=* 3.91, *P* = 0.003, *d* = 0.55, 95% CI [3.85, 19.27]). Complementarily, they had more correct trials when they entered the rooms in the same (rather than the different) sequence during encoding and retrieval (*β =* −14.3, SE *=* 5.48, *P* = 0.009, *d* = 0.40, 95% CI [−25.16, −3.52]).

Finally, to establish a direct link between context familiarization and later context memory, we examined in an exploratory analysis whether children’s engagement with context-relevant objects during familiarization predicted context error rates at retrieval. The proportion of time spent in targeted play (i.e. playing with context-specific items; see Table S1) relative to trial duration differentially predicted context error rate after sleep and wakefulness (**Fig. S2**; Condition × Targeted Play: β = 0.523, SE = 0.20, *P* = 0.018, 95% CI [0.10, 0.95]). Follow-up analyses showed that more targeted play was associated with fewer context errors after sleep (β = −0.297, SE = 0.14, *P* = 0.048, 95% CI [−0.59, −0.01]) but had no effect after wakefulness (β = 0.228, SE = 0.15, *P* = 0.139, 95% CI [−0.08, 0.54]). Thus, richer interaction with the contextual features during familiarization benefitted context memory specifically when followed by sleep. Furthermore, sleep likely did not have unspecific effects on attention and task engagement as motivation during encoding was not related to the context error rate across conditions (sleep: r = −0.054, *P* = 0.788; wake: r = −0.043, *P* = 0.830).

### Sleep enhances contextualized memory and reduces decontextualized memory

To more sensitively assess the degree to which memories were bound to their spatial context, we computed a “Contextualized Memory Index” (CMI) defined by the formula: percent correct trials / (percent context errors + percent correct trials). Thus, by normalizing the number of correct trials to the total number of correct container decisions, the CMI more specifically expresses the child’s ability to associate the location of the puppet with the correct room, i.e., the spatial context. Correspondingly, to more specifically assess the memory for the kind of container used for hiding a puppet, independently of the memory for the room context, we computed a “Decontextualized Memory Index” (DMI) defined by the formula: percent context errors / (percent context errors + percent random errors), where trials with a correct container choice but incorrect context allocation (i.e., context errors) are normalized to the total number of errors.

When accounting for the shared variance of CMI and DMI using a multivariate linear-mixed effects model including both, the CMI and DMI, we observed an increased CMI ( *β =* 0.15, SE *=* 0.07, *P* = 0.015, *d* = 0.49, 95% CI [0.01, 0.27]) and a decreased DMI (*β = −* 0.15, SE *=* 0.07, *P* = 0.017, *d* = 0.60, 95% CI [−0.30, −0.01], **Fig. 2e**) after sleep compared to wakefulness, suggesting that sleep enhances a contextualized memory, while memory for the containers is reduced after sleep. Importantly, the observed increase in CMI and decrease in DMI after sleep reflects a relative shift toward fewer context errors among all container-related decisions, consistent with more precise use of spatial context information during recall.

Effects of sleep and wakefulness on the CMI were modulated by the child’s age (**Fig. 2f**). With increasing age, the CMI decreased in the sleep condition and increased in the wake condition (*β = −*0.003, SE *=* 0.001, *P* = 0.015, 95% CI [−0.01, −0.001] for Sleep/Wake x Age interaction). Also, Sequence effects were present for the CMI, which was higher when children entered the rooms in a different sequence during encoding and retrieval (*β = −*0.17, SE *=* 0.06, *P* = 0.005, *d* = 0.57, 95% CI [−0.30, −0.05]).

### Influence of sleep vs. wakefulness on the child’s behavior during encoding

In an exploratory fashion, we examined whether sleep affected the child’s behavior already at encoding (hiding the puppets) and whether such influence could predict the formation of spatial context memory. To this end, each child’s behaviors (including interactions with the experimenter and parent) during familiarization and during encoding were scored considering 10 behavioral states (**Table S1**). Transition probabilities between the scored behaviors were computed to build directed graphs for each trial separately. **Fig. S4a** shows exemplary trial graphs.

Training a graph neural network (GNN) on 90% of the trial graphs (with a cross-validation accuracy of max. 69.97%, **Fig. S4c**) and testing it on the remaining 10% revealed above empirical chance level (49.95%) accuracy of the classification of behavioral interactions between sleep and wake conditions (65.79%, *P* = 0.043, 5000 permutations), with a model sensitivity (to correctly detect sleep trials) of 73% and a specificity (to correctly detecting wake trials) of 67%. Importantly, the GNN could not differentiate graphs between conditions at context familiarization prior to sleep or wakefulness (test accuracy: 50.00%, *P* = 0.739, 5000 permutations). Graph explanations used to determine the specific behaviors contributing to successful GNN classifications at encoding revealed a major role for transitions from “looking into the container” to “container play” in most trials (**Fig. S4d**). For those trials, the occurrence of such transitions at encoding was higher after post-familiarization wakefulness compared to sleep (*β =* 0.55, SE *=* 0.17, *P* = 0.001, 95% CI [0.21, 0.88]), and predicted memory performance: In correct trials after wakefulness, children had switched more often from “looking into the container” to “container play” compared to sleep (*β =* −0.55, SE *=* 0.17, *P* = 0.001, 95% CI [−0.88, −0.21]) and compared to trials where they committed context errors (*β =* 0.61, SE *=* 0.23, *P* = 0.020, 95% CI [0.08, 1.14]) and random errors (*β =* 0.91, SE *=* 0.30, *P* = 0.006, 95% CI [0.22, 1.61]; **Fig. 3**). Similarly, only in the wake, but not in the sleep condition, higher transition probabilities from “looking into the container” to “container play” at encoding predicted diminished DMI values (*β =* 2.16, SE *=* 0.04, *P* = 0.039, 95% CI [0.11, 4.20]) and a trend for better CMI values (*β =* −1.46, SE *=* 0.08, *P* = 0.062, 95% CI [−2.99, 0.07]).

**Figure 3.**
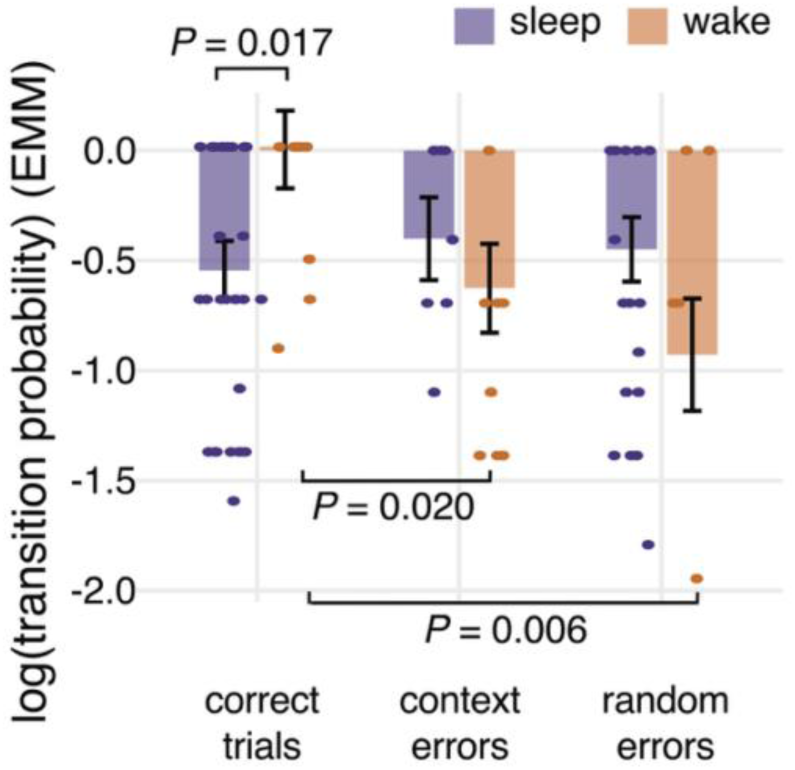
Encoding behavior associated with memory performance. Log-transformed transition probabilities from “looking into container” (“0” in **Fig. S4a**) to “container play” (“1”) for trials identified by graph explanations, separately for correct trials, trials with context errors and random errors (means ± SEM and significant differences are indicated)

### Spindles and SO-spindle phase-amplitude coupling predict contextualized memory

Based on previous findings in children^56^, we focused our analyses of EEG sleep recordings on potential contributions of spindle and SO events to the formation of spatial context representations (see **Table S2** for sleep stage architecture). Spindles and SOs were detected in all recorded channels (C3, Cz, C4) using established algorithms^39,57^ for computing spindle and SO amplitude, density, and duration, as well as SO slope (see **Table S3,** for spindle and SO characteristics). Spindle amplitude (*F*_2,64_ = 6.921; *P* = 0.001, η² = 0.18, 95% CI [0.05, 1.00]) and SO amplitude (*F*_2,64_ = 11.213; *P* < 0.001, η² = 0.26, 95% CI [0.11, 1.00]) were higher at Cz than C3 (*t*_21_ = −2.636 and −3.567; *P* < 0.005, for spindles and SOs, respectively) and C4 (*t*_21_ = −2.637 and −3.080; *P* < 0.005).

Regression models (including channel and its interaction with amplitude as covariate) revealed an association of spindle density with fewer context errors (*β* = −7.19, SE = 2.98, *P* = 0.017, r_standard_ = −0.38, 95% CI [−13.06 – −1.32]). No other spindle or SO parameters were associated with the percentage of context errors (all other spindle *P* > 0.322; SOs, all *P* > 0.324). Increased spindle density also predicted a higher CMI (*β* = 0.11, SE = 0.04, *P* = 0.012, r_standard_ = 0.38, 95% CI [0.03 – 0.20], **Fig. 4a**). Again, no other spindle or SO parameter was predictive of the CMI (all other spindle *P* > 0.121; SO *P* > 0.133). Thus, only spindle density, but not SOs per se, appear to contribute to the formation of context memory. In adults and older children, memory consolidation is known to particularly benefit from spindles co-occurring with SOs, with the 12-16 Hz spindle events frequently occurring in the down-to-up-state transition and during the upstate of the SO event ^8,39,58,59^. Our analyses of co-occurring SO-spindles confirmed that spindles tended to occur during the down-to-up state transition of the SO, also in our toddlers (**Fig. 4**). Generally, the percent of spindles co-occurring with SOs was above chance level at all channels (C3: *t*_22_ = 10.798, *P* < 0.001, *d* = 3.003, 95% CI [−2.16 – 3.85], Cz: *t*_21_ = 13.547, *P* < 0.001, *d* = 4.054, 95% CI [3.02 – 5.09], C4: *t*_21_ = 11.677, *P* < 0.001, *d* = 3.330, 95% CI [2.42 – 4.24]). Peri-event time histograms around the negative half-wave peak of the SO showed the characteristic SO-spindle coupling pattern with enhanced spindle occurrence rates between 400-600 ms after the negative SO half-wave peak. This coupling pattern, however, reached significance only at Cz (**Fig. 4b**). Analyses of phase-amplitude coupling, on the group level, showed significant SO-spindle phase-amplitude coupling in all channels (C3: z = 9.335. *P* < 0.001; Cz: z = 12.0640, *P* < 0.001; C4: z = 10.03, *P* < 0.001), with a trend for strongest coupling over Cz (*F*_2,64_ = 2.718, *P* = 0.074, η² = 0.08, 95% CI [0.00, 1.00]). At the individual level, no robust phase-locking of spindle amplitudes to the SO upstate was present at Cz in 3 participants and at C3 and C4 in 7 participants. Given that phase-locking was most robust at Cz, in further analyses examining the relevance of SO-spindle coupling for forming contextualized spatial memories, we focused on recordings at Cz.

**Figure 4.**
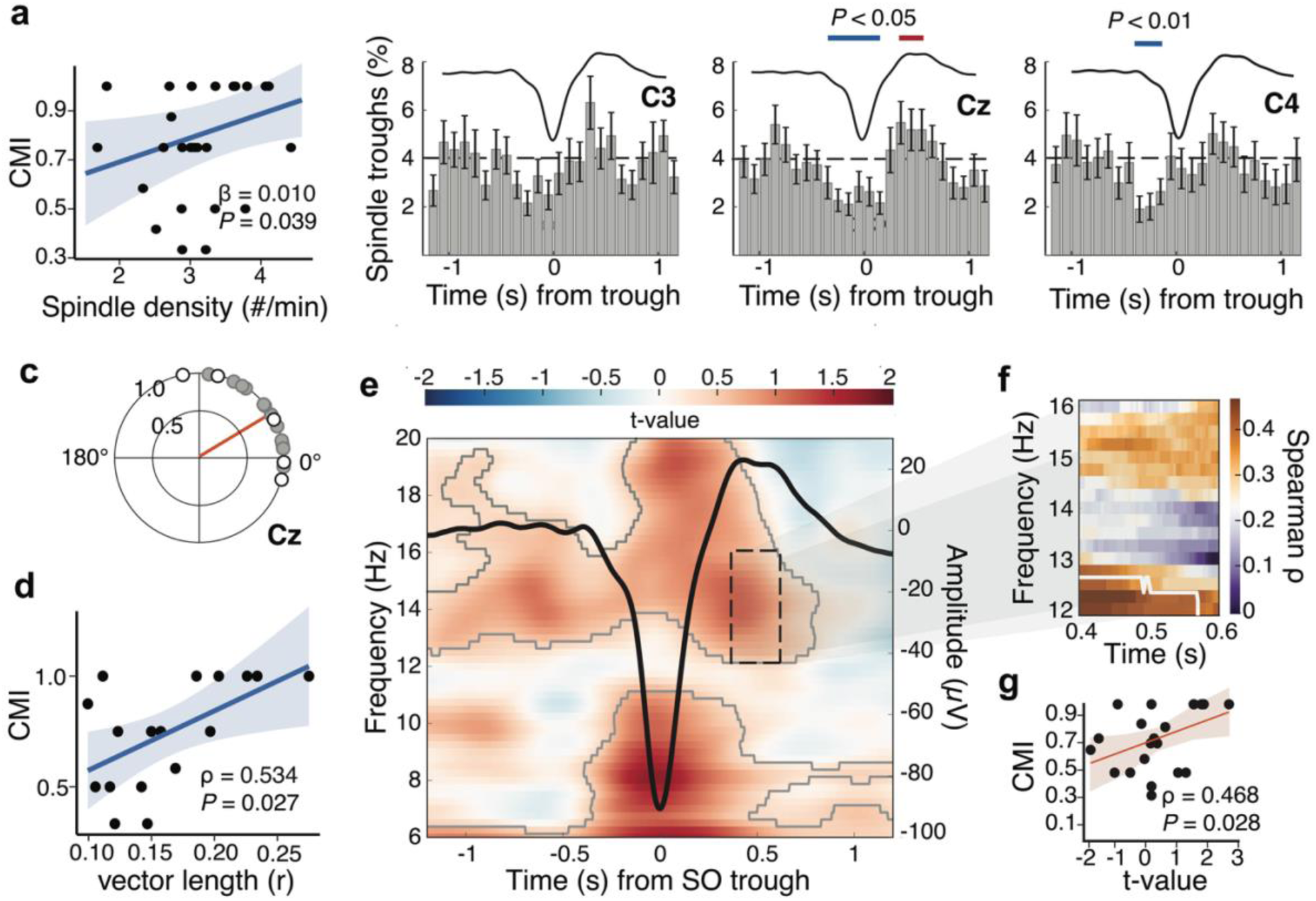
SO-spindle coupling correlates with contextualized memory. **a**, Partial robust correlation between spindle density and CMI scores (indicated by predicted value of the CMI for spindle density; raw values overlaid, significance for regression coefficient, N = 25). **b,** Peri-event time histograms (PETHs) for occurrence of spindle (troughs) during the SO, time-locked to SO negative half-wave peak (trough, 0 s). Error bars indicate standard errors per bin. Black dashed lines indicate baseline level derived from surrogate data. Average SOs are superimposed. Blue and red lines indicate significant negative and positive clusters, respectively. **c,** Preferred SO-phase of spindle amplitude maxima at Cz across participants. Vector direction indicates the circular group mean and vector lengths the consistency in coupling. Overlaid dots show the preferred phase for each participant. Filled dots indicate participants with a significant non-uniform phase-amplitude coupling (*P* < 0.05, N = 22) **d,** Associations of preferred phase (see panel **c**) with CMI scores (effect size and significance are indicated for Spearman correlation, N = 17). **e,** Time-frequency representations (TFRs) of co-occurring SO-spindle events (at Cz). Color code refers to difference in power with reference to isolated SO events lacking a co-occurring spindle, expressed as t-value. Grey lines outline significant positive cluster (bootstrapped, cluster-corrected one-sample t-tests against 0). **f,** Spearman correlation coefficients (color coded) between CMI scores and the t-values for each frequency bin in the 400-600 ms square window indicated in panel **e.** White line outlines significant cluster (bootstrapped, cluster-corrected). **g,** Pixel of panel **f** with maximum positive correlation (N = 22).

Generally, SO-spindle co-occurrence rates were not associated with CMI (ρ = −0.104, *P* = 0.644), and so was the phase-amplitude coupling for the entire 12-16 Hz range (ρ = 0.203, *P* = 0.363). However, we found a significant positive association between SO-spindle phase-amplitude coupling and the CMI score selectively for the slower 12-13 Hz spindle frequency range (ρ = 0.534, *P* = 0.027, **Fig. 4d**), consistent with the observation that in younger children slower spindle-frequency ranges are more prominent^39,56^. For the faster spindle frequency bands (13-14 Hz, 14-15 Hz, 15-16 Hz) the association with the CMI remained non-significant (all *P* > 0.111). Importantly, phase-amplitude coupling at 12-13 Hz was directly associated to the CMI value (ADE = 2.65, *P* = 0.011), with no evidence that this association was mediated through the effect of spindle density described above (ACME = 0.03, *P* =Pyto0.936).

This finding of a spindle-frequency specific link between SO-spindle phase-amplitude coupling and the CMI was confirmed in a time-frequency resolved analysis of 12-16 Hz spindle power in the designated 400-600 ms window (following the SO negative half-wave peak). As expected, spindle power was distinctly higher during coupled compared to non-coupled SO events lacking a co-occurring spindle (*t*_sum_ = 10,991, cluster-corrected *P* = 0.028, **Fig. 4e**). Correlating the power difference between coupled and non-coupled SO events with the CMI, separately for each time-frequency point in the 400-600 ms window between 12-16 Hz, we found a significant positive relationship between CMI and SO-spindle coupling power at 12-13 Hz (ρ = 0.422, cluster-corrected *P* = 0.033, **Fig. 4f**), with a maximum correlation of ρ = 0.468 (*P* = 0.028, **Fig. 4g**).

In exploratory analyses, we additionally assessed the correlations between major parameters of macro-sleep architecture and measures of context memory. None of these parameters were significantly associated with percentages of context errors (all uncorrected *P* > 0.236) or CMI scores (all uncorrected *P* > 0.304, **Table S4**).

## Discussion

We assessed the effects of sleep on the formation of abstracted context memory for two experimental rooms in toddlers aged 2-3 years, an age when episodic and spatial memory capabilities are beginning to emerge. Our main finding indicates that a nap following familiarization with the two rooms indeed enhanced memory for these contexts. This was evidenced by fewer context errors and a higher context discrimination index (CMI) in a hide-and-seek task conducted after sleep, compared to performance after a wake period following familiarization. The post-familiarization nap already changed the toddler’s behavior during the encoding (“hide”) phase of the hide-and-seek task such that, unlike after wakefulness, the toddlers in the sleep condition did not need to transit from opening to playing with the containers for successfully discriminating the spatial contexts. The enhancing effect of sleep on signs of context memory was associated with increased spindle activity and SO-spindle coupling during NonREM sleep. Together, these findings indicate that sleep supports, already at this early stage of cognitive development, the formation of context memory that is abstracted from the originally experienced familiarization episode and subsequently used as contextual reference for forming novel object-context associations, guiding behavior in a hide-and-seek task.

Our finding of enhanced signs of context memory after a nap period of approximately 90 min in toddlers aligns with studies in children demonstrating that rudimentary capabilities to form context memory emerge early during development when the hippocampus – assumed to mediate context memory – still exhibits structural and functional immaturity^24,32,60,61^. We adapted our hide-and-seek task paradigm from a study by Newcombe et al.^46^, which aimed at assessing children’s ability to bind incidental features of the (room) contexts with the object (i.e., the container with the hidden toy). These authors reported that even 15-month-olds exhibited a fragile ability to discriminate between room contexts during retrieval, an ability that improved gradually across childhood up to 6 years of age. A rather early onset of capabilities to form context representations is likewise suggested by studies of juvenile rodents^62^. However, it should be noted that these studies used rather short retention intervals, typically not extending over more than a few minutes. Juvenile rats tested on object-in-context and contextual fear conditioning paradigms on postnatal day 17 (roughly comparable with human infancy) showed context memory at a test after 5 min but not after 24 hours^63^. Against this backdrop, our findings are the first to demonstrate signs of more persistent context memory in human toddlers, lasting for more than 1.5 hours.

Importantly, toddlers formed persistent context memories only if they slept following familiarization. Context errors, i.e. allocating a correct container to the incorrect room, were significantly below chance level only in the sleep condition, with the sleep benefit being particularly pronounced among the younger children in our sample. Reduced tiredness is unlikely to explain the sleep effect, considering its specificity: performance regarding the percentage of correct trials and random errors was comparable between the sleep and wake conditions. Additionally, motivation ratings from the experimenter in each trial did not differ between conditions. This specific memory enhancement after sleep therefore highlights sleep’s critical role for consolidating memories into a long-term store — here, for newly acquired context representations — which appears to be similarly important during childhood as in adulthood^3,22,32,33^.

Our task design differed from classical object-in-context association paradigms such that sleep versus wakefulness was manipulated *after* toddlers were familiarized with two room contexts (subsequently used in the hide-and-seek task to form novel object-in-context associations). Introducing sleep after familiarization and not after encoding (i.e., hiding the toy) allowed us to isolate the effects of sleep on pure context memory, rather than specific context-event binding. In line with our hypothesis, we found that children were less likely to confuse the correct container holding the hidden puppets between the two contexts when they had slept after context familiarization. Notably, sleep did not affect other memory performance measures, such as the percentage of correct trials or random errors, both of which conflate the assessment of contextual memory (for the rooms) with aspects of immediate episodic memory formed during the hide-and-seek task, namely the memory for the container to be associated with one of the room contexts. A specific effect of sleep on contextual representations is further indicated by our analyses the “Context Memory Index” and the “Decontextualized Memory Index”, i.e. indices that aimed to more purely separate context from object aspects (container with the puppet) in memory performance. Sleep, in these analyses, was found to increase the Context Memory Index, with this effect being even at the expense of decontextualized memory. In combination, these results corroborate the view that the toddler’s sleep is quite effective in forming distinct spatial context representations^55^.

Interestingly, sleep benefits for memory contextualization were most pronounced in younger toddlers (roughly 24-30 months) considering that sleep-related reduction in context errors was attenuated with increasing age. This age range may reflect an early developmental window in which wake-based maintenance of distinct context representations is particularly fragile, rendering toddlers prone to context confusions unless sleep intervenes. This account aligns with developmental evidence that flexible contextual what–where binding shows a marked inflection around the second year but continues to refine over subsequent years, and with proposals that the transition out of naps reflects increasing hippocampal capacity to retain newly encoded information across wake^22,49,64^. In addition, age-related improvements in attentional control and emerging executive strategies may allow older toddlers to disambiguate contexts during wake by relying on online cues, reducing context errors even without sleep-dependent abstraction^49^. Together, these findings suggest that sleep confers its strongest benefit for context memory when wake-based retention of contextual information is still fragile.

Consolidation during sleep is characterized by several EEG features, including SOs, spindles, and hippocampal ripples occurring during NonREM sleep, which are thought to mediate an underlying dialogue in memory processing between hippocampal and thalamo-neocortical systems^3,32,65^. Spindle activity, in particular, has been linked to memory consolidation also during early development, starting at three months of age^19^. Perhaps even more relevant than spindle occurrence per se is the temporally precise synchronization of spindles with the SO ^35,40,66^. While an enhanced co-occurrence of SOs and spindles has been observed already in 6-month-olds, significant phase-amplitude coupling between SOs and spindles emerges somewhat later, around 14 months of age^39^, and continues to increase throughout childhood and adolescence^38,56^. Notably, no study to date has linked this coupling pattern to memory consolidation in children as young as 2-3 years. Against this backdrop, our findings are the first to show that beyond spindle density^15,35^, also the precise phase-coupling of spindle amplitudes to the SO-upstate predicts better contextual memory. The coupling mainly pertained to the slow 12-13 Hz range of spindle activity which might reflect a general age-related prominence of lower spindle frequencies^67–69^. However, spindle frequency also shows topographical variations, as does SO-spindle phase-amplitude coupling^36,67,68^. Hence, a limitation of our study is that, for practical reasons, EEG recordings were restricted to central electrode sites. Future research needs to address the question whether consolidation of contextual information is particularly linked to region-specific SO–spindle coupling, which is often most prominent over prefrontal cortical areas^14,40^.

Spindles, especially when nesting in the SO upstate, are associated with enhanced synaptic plasticity in neocortical circuitry^13,70,71^. In the mature brain, coupled SO-spindle events are associated with the occurrence of ripples and memory replay in hippocampal networks^10,72^. Signs of hippocampal memory replay activity have also been observed in sleeping human toddlers, with increased hippocampal reactivations linked to better memory performance ^73^. In conjunction with this evidence, our findings of phase-amplitude SO-spindle coupling predicting enhanced context memory in the toddlers would agree with the concept of an active systems consolidation assuming that the consolidation process during sleep originates from hippocampal replay of contextual information during NonREM sleep ^3,7,74^.

Alternatively, the sleep-dependent enhancement of context memory which we observe in our toddlers may be a consequence of a consolidation process that is restricted to the thalamo-cortical system with negligible inputs from hippocampal memory replay-related activity. This view of a predominant consolidation within the thalamo-cortical system is not unlikely given the evidence for structural and functional immaturity of the hippocampus at this early age^32,36^. Structural MRI shows continued volumetric growth of the hippocampus across the first two years of life^42^ and protracted, subfield-specific maturation well into childhood and adolescence, with CA1, dentate gyrus and subiculum volumes continuing to change across early and middle childhood and relating to age-dependent improvements in memory^75,76^. Diffusion-based measures similarly indicate ongoing refinement of hippocampal microstructure between 4–8 years that is correlated with better episodic memory^77^. Together, these findings support the view that, at 2–3 years, the hippocampal system is still immature.

Moreover, although in episodic memory research the encoding of (spatial) context is commonly allocated to the hippocampus^78^, the hippocampus may not be the only structure serving this function^79,80^. In rodent studies, lesions to the hippocampus had only minor effects on the freezing response in context fear conditioning. Noteworthy are experiments in adolescent rats using a “context pre-exposure facilitation” procedure which shares similarities with the present task paradigm in that the rats are familiarized with the context itself on the day before acute context-shock conditioning^81,82^. Inactivating the medial prefrontal cortex during context pre-exposure impaired contextual fear conditioning in the same way as did the inactivation of the ventral hippocampus. While these studies did not examine the role of sleep, they show that context information during familiarization is encoded into multiple traces, one residing in the hippocampus and another one involving the medial prefrontal cortex.

Accordingly, the sleep-dependent enhancement of context memory which we observe in our toddlers may alternatively be a consequence of a consolidation process that is restricted to the thalamo-cortical system with negligible hippocampal inputs. It is important to note that in the present study we did not collect direct measures of hippocampal structure or activity. Thus, the question whether sleep-dependent consolidation during early life involves, like in adults, a dialogue between hippocampus and neocortex, or represents a process restricted to thalamo-cortical circuits, remains to be resolved in future research. Given the distinct dependency of the memory effect of sleep on age – apparent even in the present rather homogenous sample of 2-3-year-olds – it seems promising to include in such research even younger children in the infant age.

Whatever the hippocampal contribution to the consolidation process is, memory processing within the thalamo-cortical networks during sleep is thought to support the formation of more generalized and abstract representations during early development^83–85^. It is worth stressing here that rather than directly testing memory for context experienced during the familiarization period before sleep, our task indeed probed an abstracted form of context memory – decoupled from the original familiarization experience –that, in the hide-and-seek task should be used for forming novel object-context associations. Our behavioral analyses confirmed that sleep after familiarization did not only improve retrieval of contextualized memory but shaped behavior already during the encoding (hiding) phase of the task. After sleep, children did not rely on an interaction with the containers to remember the puppet locations, indicating the use of a representation of the context rather than a puppet-container association to perform the memory task. Such transfer function, i.e., to use memorized contextual information for coping with novel task situations is commonly ascribed to neocortical, specifically prefrontal cortical regions^86,87^, which, notably, is also functionally immature in toddlers^49^.

### Limitations and future directions

Despite the advancement in understanding sleep effects on spatial context memory formation, several limitations of the present study should be acknowledged. One major limitation concerns the interpretation of our findings with respect to sleep effects on representations of the contexts experienced during the familiarization phase versus effect on encoding of the hide-and-seek task. In our paradigm, the familiarization phase establishing contexts preceded the sleep/wake manipulation, whereas the encoding of object–location associations and their retrieval in the hide-and-seek task occurred thereafter. Control analyses revealed a sleep-specific relation between behavioral indicators of familiarization strength (targeted play) and later context errors while controlling for behavioral sequences during encoding affected by sleep (i.e. looking into container → container play). Together with the absence of condition differences in motivation at encoding, these findings are consistent with the view that sleep strengthened an underlying representation of the spatial contexts independent of its effects on encoding processes. Indeed, non-specific effects of sleep, e.g., increasing alertness are highly unlikely to express themselves in a specific enhancement of context memory but would have been expected to similarly enhance object memory (i.e., to decrease random errors^88^. At the same time, we cannot fully exclude that sleep also modulated subsequent encoding processes. A problem that is difficult to resolve by appropriate control conditions in the context of the present task paradigm and may be addressed in future work.

Additionally, sleep and wake conditions were not fully matched for time-of-day. While all sleep sessions occurred during the toddlers’ habitual midday nap, wake sessions took place either in the morning or in the afternoon. Consequently, circadian phase and time-of-day–dependent fluctuations in arousal cannot be entirely ruled out as contributing factors. However, several observations argue against a primary role of such effects. First, prior wake duration did not predict any of the memory measures when included as a covariate, nor did it interact with the Sleep/Wake condition. Second, within the wake condition, memory performance did not differ between children tested in the morning versus the afternoon. Finally, as mentioned above, the observed sleep effects were selective to measures of contextual memory, whereas overall accuracy and random error rates were unaffected, which would be unexpected if general alertness or fatigue were driving the results. Although a fully time-matched design would be preferrable to rule out circadian effects, it is not possible without sleep depriving habitually napping children, leading to confounding effects of fatigue during the memory task^89^.

Another methodological consideration concerns differences in the total duration of the break period between the sleep and wake conditions. Although naps were not terminated prematurely and could therefore exceed 120 minutes, a ∼30-min buffer period was included after both sleep and wake retention intervals to equate the effective delay before memory testing and to minimize sleep inertia effects. Additional control analyses confirmed that retention interval duration was not significantly different between conditions, did not predict memory performance, and did not account for the observed sleep-related reduction in context errors and increase in contextualized memory.Future studies may consider strictly matching retention intervals across conditions to further address potential time-dependent decay of memory representations. However, such an approach would come at the cost of either partially sleep-depriving participants or introducing systematic order effects if the sleep condition were always scheduled before the wake condition.

It should be noted that our sample comprised toddlers aged 2–3 years, i.e., slightly beyond the 18–24-month period that has been proposed as particularly critical for the onset of hippocampal-dependent memory. While converging evidence indicates substantial continued hippocampal and episodic memory development between 2 and 5 years, our findings therefore speak specifically to an early toddler stage, in which context memory is present but still immature, rather than to the very first emergence of hippocampal-dependent memory. Whether similar sleep-related benefits for contextual memory and similar thalamo-cortical mechanisms can already be observed in younger infants (< 25 months) remains an open question. Future work adapting the current paradigm to earlier developmental stages will be essential to determine how early in life the sleep-dependent consolidation of spatial context representations emerges and how it changes across the extended trajectory of hippocampal maturation.

A notable aspect of our findings is the modulation of sleep effects by age within a relatively narrow 25-34-month range. Although our two-room paradigm is known to be sensitive to age-related differences in context memory^46^, our restricted age range warrant a cautious interpretation of these effects. To capture the narrow age range more precisely, we expressed age in days rather than months. Although the modulating effect of Age remained, significant when age was expressed in a more fine-grained way, in days rather than months, it was not significant in follow-up comparisons and individual datapoints showed large variability. We therefore view the present age effects as suggestive rather than definitive, consistent with the idea that sleep-related benefits for context memory may be particularly pronounced at the younger end of the toddler period, but in need of replication in larger samples and with broader age ranges.

## Methods

### Participants

Initially, 41 toddlers were recruited for participation in the study. Five children dropped out during data acquisition, four after the screening session and one during the first experimental condition. In addition, eight children were excluded post hoc because they did not complete any trials in at least one condition without experimenter assistance, either due to insufficient cooperation (N = 7) or illness (N = 1). Thus, the final sample consisted of 28 toddlers (29.46 ± 2.57 months; range: 25–34 months; 12 female). Inclusion criteria, assessed via a structured telephone screening, required full-term birth, good health, no regular medication, age-appropriate development, and routine midday naps. Cognitive development was assessed during an in-person screening session using the Bayley Scales of Infant and Toddler Development, Second Edition (Bayley-II), yielding a Mental Development Index (MDI) of 102.55 ± 7.51 (range: 82–114). Upon successful completion of the study parents received monetary compensation, and children received a small toy. The study was approved by the ethics committee of the university clinic in Tübingen, and written informed consent was obtained from all caregivers beforehand.

### Design and General Procedure

The experiments were conducted as a randomized within-subject cross-over design. After a screening session, each toddler was tested on two experimental conditions, a sleep and a wake condition (**Fig. 1**). During the screening session, children were familiarized with the experimenter and the laboratory setting, performed the Bayley II Developmental Test, and were introduced to a portable dummy EEG, which the parent took home and equipped the child with during one nap in the week leading up to the first test condition for adaptation purposes.

The toddler’s two experimental conditions were at least two weeks apart and in two different locations but followed an identical procedure. Each condition consisted of three periods: a familiarization period to establish context representations for the rooms, a retention period filled with a nap or wakefulness, respectively, and a spatial memory task used for probing the context representations. During the familiarization period, participants were playfully introduced to two rooms, which were each decorated according to a theme (**Fig. S3**). During the following retention period, participants either slept for approx. 90 min (92.94 ± 5.81 min, range: 27.00 – 159.50 min) while we recorded polysomnography or they stayed awake for 90 min. In the sleep condition, children were not awakened prematurely, allowing the retention period to exceed two hours. In the wake condition, participants went with their parents for a walk and could engage in quiet activities. Following the ∼90-min retention interval, both sleep and wake conditions included an additional ∼30-min period before the memory task, either to allow recovery from sleep inertia (sleep condition) or to ensure comparable transition into the task (wake condition), respectively. The spatial memory task was a hide-and-seek game comprising 4 trials. Each trial included a short 5-min delay between hiding and searching the puppet. Parents were always passively present during the test conditions.

Because of the young age of the participants, we chose that each child completed all sessions with the same experimenter to ensure familiarity and reduce stress-related variability. Consequently, the experimenter was not blinded to the child’s condition. To minimize expectancy effects, all instructions and prompts during familiarization, encoding, and retrieval followed a strictly standardized script that was applied identically across sleep and wake conditions. The experimenter was trained to use only neutral, predefined verbal prompts and to refrain from providing corrective feedback or hints. Parents were present throughout the task but were instructed to remain passive and not to interact with the child.

The daytimes for the experimental sessions were adjusted to the child’s individual chronotype^90^. Since we did not conduct a sleep deprivation study, wake appointments were either in the morning (*N = 15*) or in the afternoon (*N* = 13) depending on the activity levels of the child, while all sleep appointments were during midday.

### Spatial Contexts

Experiments were conducted in four distinct rooms in two locations in the city of Tübingen (lab A vs. lab B). Each room served as a play environment themed with specific visual and interactive cues. Lab A featured the play kitchen and teddy clinic contexts, while Lab B hosted the zoo and train station. Each room included four containers that served as potential hiding spots during the memory task. Containers differed across labs A and B to prevent generalization. Context-specific props were positioned atop or adjacent to the containers to facilitate contextual encoding (see **Fig.S3** for each room’s layout).

During the hide-and-seek phase, one hand puppet per context was repeatedly hidden in and retrieved from the containers by the child. The puppets were not tied to specific contexts. For each test condition, two puppets were randomly selected out of eight available ones, ensuring randomized puppet assignments both between and within participants.

### Familiarization

Children explored each context for 15 minutes under guided play using a standardized script. The experimenter joined the child’s play after a minute of free exploration, introducing room elements and prompting engagement with containers via contextualized narratives (e.g., “Let’s cook an egg here—is there something inside?”). Each container was opened by the child to ensure encoding of its location and identity. Familiarization was documented using structured protocols. Participants spent 15 minutes in each context.

### Spatial Memory Task – Hide-and-Seek Game

To assess the influence of sleep on the context memories encoded during the familiarization period, the children’s capability to form context-specific object-location associations was tested in the familiarized contexts, using a hide-and-seek game. The game was following a procedure by Newcombe et al^46^ including several modifications.

During encoding, children were instructed to hide one puppet in a specified container in each of the two familiarized contexts. The sequence in which the child entered the contexts during encoding was counterbalanced across trials and participants. In contrast to the original paradigm by Newcombe et al^46^, children were animated to actively place the puppets in the target container according to the instructions of the experimenter. The designated target container differed between the two contexts within each trial to prevent participants from relying on a single hiding location across rooms. Within a trial, the hiding locations were spatially opposite for the two rooms and across trials. The sequence of target containers was counterbalanced between participants. The encoding phase followed a standardized procedure. First, all four containers were inspected with the child to confirm that they were empty (“Did you see? All the hiding places are empty?”). The experimenter then provided specific instructions on where to hide the puppet, ensuring that children actively placed the object themselves to enhance episodic memory formation. Before leaving the room, children were reminded to remember where they had hidden the puppet. The same procedure was repeated for the second puppet in the second context.

Encoding was followed by a 5-min delay to avoid context interference effects. During this time, the toddlers put stickers in a sticker book in a third, separate room. Then, they rang a bell indicating the upcoming retrieval (seek) phase.

During the retrieval phase, children were asked to recall the puppet locations and search for the hidden puppets in the two contexts. Retrieval began either in the same context as encoding or in the other context, balanced across trials. Upon entering the room, children were prompted to find the puppet. If the puppet was not located independently by the child within 2.5 min, the experimenter opened the correct container with the puppet to prevent frustration; however, performance outcomes were defined based on the child’s first container choice prior to any intervention. Since the potential hiding locations were identical across both contexts, successful retrieval depended on the child’s ability to recall the specific context in which a puppet had been hidden during encoding.

### Video Recordings and Scoring

All familiarization and encoding phases of the spatial memory task were video recorded using Axis M1065-LW IP cameras. Recordings were analyzed with BORIS software (v8.21.10). One participant was excluded due to technical failure. Behavioral scoring focused on a predefined set of 10 target behaviors (**Table S1**), capturing child, parent, and experimenter actions relevant to task engagement and encoding. The coding scheme was developed iteratively based on five independent pilot datasets, following established best practices for behavioral video coding (Datavyu User Guide, PLAY Project). An initial coding scheme was defined on one dataset and subsequently refined across the remaining pilot datasets through repeated rounds of error checking, inter-rater reliability assessment, and descriptive analyses, resulting in the final set of behavioral states.

All behaviors were defined using explicit coding criteria summarized in a detailed coding manual specifying behavioral definitions, temporal boundaries, and actor assignments. Behaviors were coded as either point events (occurring at a single time point) or state events (with defined onset and offset). From point events, we derived the total duration of each encoding trial, i.e. the time from “room entrance” to “room exit”. Due to the time-intensive nature of manual video coding, all videos were scored by a single primary coder in multiple passes who was blind to the experimental conditions. Inter-rater reliability was assessed on a randomly selected subset of 20% of the videos using an independent reliability coder, following a first-coder/reliability-coder approach. Agreement was quantified using Cohen’s κ computed over 2-s time windows, yielding substantial inter-rater reliability (κ = 0.73).

Among the target behaviors, the rater coded a state event termed “targeted play” as a marker of context familiarization^91^. Targeted play was defined as periods during which the child actively engaged with objects that were unique to the specific room (e.g. zoo animals, doctor’s kit, kitchen utensils or train tracks), excluding items that were identical across contexts (containers, furniture, door handles, etc.). For the familiarization in each room, we computed the percentage of time spent in targeted play relative to the total familiarization duration. This measure was used to test the relevance of context familiarization for the two conditions. The rater also coded any verbal or non-verbal actions by the experimenter or parent that could potentially cue the child. Trials containing non-allowed prompting were excluded from analysis.

Motivation during encoding of the hide-and-seek task was rated from the video recordings on a 0–10 scale adapted from ^92^, capturing mastery motivation (persistence, competence), strategy use (help-seeking), and emotion expression (e.g. positive affect, interest/arousal, frustration). Two independent raters scored each trial; ratings were averaged across raters and across trials within each condition. Trials where both raters scored 0 motivation were excluded from analysis. Motivation scores were used in control analyses to test whether differences in context memory could be explained by general engagement or alertness at encoding.

### Sleep-EEG Recordings

A portable polysomnography device (SOMNOscreen™ plus, SOMNOmedics) was used for the sleep-EEG recordings (sampling rate: 256 Hz). The polysomnographic recording included electroencephalography (EEG), electrooculography (EOG), and electromyography (EMG). EEG electrodes were individually placed according to the 10–20 system and recorded from C3, C4, M1 and M2 (Cz as reference, FPz as Ground). Additional electrodes included two EOG electrodes (placed above the right eyebrow and below the left eye), and two EMG electrodes positioned on the chin. EEG data was acquired for all but three participants due to insufficient cooperation during electrode placement. Thus, respective sleep EEG analyses included recordings from N = 25 toddlers.

### Behavioral Outcome Measures

Primary behavioral outcome measures were the numbers of correct trials, context errors, and random errors, all expressed as percentages with the individual total number of trials set to 100 % (see also **Fig. 1c**).

To dissociate different aspects of recall behavior, we derived two complementary indices from the same response distribution. The Contextualized Memory Index (CMI) is defined as:

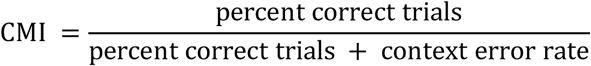

The CMI reflects the proportion of successful context–object associations out of all context-relevant responses and quantifies the child’s ability to bind object memory to its spatial context. Conversely, to isolate memory for container identity independent of spatial context, we computed a Decontextualized Memory Index (DMI) as:

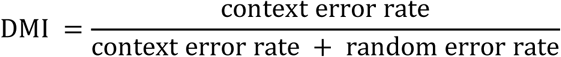

By excluding trials with accurate context placement, the DMI specifically indexes container memory stripped from context-binding influences.

Both indices share the context error rate and are therefore statistically dependent. Experimental effects on CMI and DMI are therefore interpreted as differential parametrizations of the same recall behavior rather than as evidence for separable or competing memory systems.

### Graph Neural Network Analysis

The video-coded behavior during familiarization and encoding yields rich temporal sequences of discrete states (e.g., looking into a container, playing with context-relevant objects, seeking help from the parent or experimenter). Rather than reducing these sequences to a few summary counts or durations, we modeled them as behavioral graphs, in which nodes represent behaviors and directed, weighted edges represent transition probabilities between behaviors within a trial. This graph-based representation preserves information about both which behaviors occur and how they are chained over time. We then used a graph neural network (GNN) as an exploratory, hypothesis-generating tool to ask (i) whether the structure of encoding behavior contains information that discriminates sleep from wake trials and (ii) which specific behavioral transitions are most informative for that discrimination and for subsequent memory performance. In this way, the GNN analysis complements our conventional behavioral measures by capturing the organization of children’s interactions with the context and containers, rather than only their overall frequency or duration, and provides candidate behavioral signatures of how post-familiarization sleep may alter encoding strategies

The GNN analysis was performed in Python 3.8, using PyTorch (v2.8.0), PyTorch Geometric (v2.5), NumPy, scikit-learn, pandas, Matplotlib, NetworkX, and Seaborn. Random seeds were fixed across libraries for reproducibility. We computed transition matrices for each encoding trial and familiarization phase in the sleep and wake conditions, expressing the transition probabilities between all target behaviors for a given trial (**Table S1**). The transition matrices served as adjacency matrices that could be converted into graph objects. These trial graphs, containing transition probabilities as edge features as well as trial metadata as node and graph features, were then subjected to a self-constructed GNN for classification. To identify the nodes contributing most to the GNN classification, we used the GNNExplainer algorithm. See Supplementary Methods for analysis details.

### EEG Analyses

Data were preprocessed in Matlab 2024b (Mathworks Inc., Sherbom, Massachusetts) using the Sleeptrip toolbox (www.sleeptrip.org; RRID:SCR_017318). All EEG channels were re-referenced to the average mastoids (M1 and M2) and the data were band-pass filtered between 0.3 and 35 Hz (4th-order Butterworth zero-phase filter). Channels were rejected based on visual inspection of the power spectra. For the remaining channels, 30s-EEG epochs with artifacts labeled as movement arousals during sleep scoring were rejected from further analysis; for 3 participants channels Cz and C4, and for 4 participants channel C3 were excluded.

Sleep scoring was done visually by two experienced and independent scorers following standard criteria (AASM, Berry et al., 2017). Inter-rater reliability after the first round of scoring was 86.232 ± 1.238 (range: 70.900 - 95.880). All epochs on which the two raters disagreed were reviewed and matched between scorers. TST was calculated as time spent in stage 1, 2, SWS, and REM sleep. WASO was defined as the time participants were awake between sleep onset and final awakening (**Table S2**).

### Event Detection

Spindle and slow-oscillation (SO) events were detected automatically using the Sleeptrip toolbox with details on the algorithms described elsewhere^93^ and the Python implementation of FOOOF^94^. For each participant, we first estimated the individual spindle peak frequency (13.795 Hz ± 0.199 Hz, range: 12.170 – 15.650) by fitting and subtracting the aperiodic 1/f component of the N2/N3 power spectrum with a Lorentzian function, then iteratively applied Gaussian fits to the remaining periodic peaks. Spindles were detected in channels C3, Cz and C4 in the band-pass filtered signal around this personalized peak (±2 Hz, 4th-order zero-phase Butterworth, see Supplementary Methods). For each individual and channel we derived the spindle density (#/min), amplitude (µV), frequency (Hz) and duration (s). Similarly, for detected SOs (see Supplementary Methods) we obtained for each individual and channel, the SO density (#/min), amplitude (µV), slope (µV/s) and duration (s).

### SO-Spindle Events

First, we determined whether spindles and SOs co-occurred significantly above chance level by calculating for each individual and channel the percentage of NonREM sleep with SOs (total SO duration/NonREM duration × 100). If spindles are distributed equally across NonREM sleep and do not tend to occur together with an SO, their proportion of co-occurring with an SO should not significantly deviate from the proportion of NonREM sleep with SOs^30^.

Furthermore, we calculated for each trough-locked SO, the proportion of spindles whose central trough fell within ±1.2-s of the SO trough—covering an entire slow-oscillation cycle. Overlapping SO windows were resolved by retaining only the window closest to each spindle center.

To visualize co-occurrence, peri-event time histograms (PETHs) were constructed by counting spindle troughs in 100-ms bins over ±1.2 s around each SO trough, then normalizing by the total spindle count within that window. Statistical significance of the PETH structure was assessed via 1000 surrogate histograms generated by randomly shuffling bin assignments; surrogates were averaged per subject and channel to form a null distribution.

For event-locked cross-frequency coupling, we extracted artifact-free SO epochs (±3 s around each SO trough), band-pass filtered them at 0.3–1.25 Hz (6th-order Butterworth, zero-phase) and computed the instantaneous phase via the Hilbert transform. The same epochs were filtered in the average spindle band (12–16 Hz) and in 1Hz frequency bins of the spindle band, from which we derived the amplitude envelopes. To avoid edge artifacts, analyses were restricted to ±1.2 s around the SO trough. Within each trial, channel, and subject, the maximal spindle amplitude (for the average spindle band and each frequency bin), and its corresponding SO phase was extracted.

Time–frequency dynamics of SO versus baseline epochs were examined using adaptive superlets^95^. The ±3 s epochs around SO-spindle events were paired with randomly selected, artifact-free baseline segments from the same sleep stage in which no spindle occurred during the SO. A fractional adaptive superlet transform was applied between 6 and 20 Hz in 0.25-Hz steps, using an initial Morlet wavelet order of 3 and an order range from 5 to 15, yielding enhanced time–frequency resolution compared to conventional wavelet methods.

### Statistical Analysis

All statistical analyses were performed in RStudio (Version 2023.12.1, R Core Team) and in Matlab 2024b. Before applying linear models, model residuals were tested for normality with the Shapiro-Wilk test and, whenever necessary, for homoscedasticity with the Breusch-Pagan test. Model parsimony was determined using the Akaike Information Criterion, Bayesian Information Criterion and log-likelihood with a backward-elimination procedure. The model with the highest trade-off between complexity and model fit is reported. Initial control analyses between sleep and wake conditions were performed using paired t-tests and Pearson correlations. Given the strong directional hypotheses for the effects of sleep on percent correct trials, random errors and context errors against chance level, one-tailed tests were applied. All other tests were two-tailed. For all tests a *P*-value of 0.05 was considered significant after Holm-Bonferroni multiple-comparison correction. Effect sizes were determined using the *effectsize* package.

Percentages of correct trials, context errors, and random errors were analyzed separately using linear-mixed-effects models (LMMs) implemented with *lme4*. For each outcome, we fitted a LMM with fixed effects for Sleep/Wake, Age (in days) given the narrow age range of our sample, Sequence (same vs. different), and their interactions, and a random intercept for each participant. In additional control analyses, Retention Interval (in min) and prior Wake Duration (in min) were included as confound predictors. Most parsimonious fixed-effect model formulas for context error rate were Sleep/Wake * Age + Sequence, while models for correct trials and random errors were Sleep/Wake + Age + Sequence.

The selectivity of the sleep effect on context errors, but not on performance accuracy, was tested with an LMM implemented with *nlme*, combining performance accuracy and context error rate into one variable (Outcome) which was treated as a factor in the model (**Table S5**). We included fixed effects for Outcome (context errors vs. correct trials), Sleep/Wake, Age, Sequence, and their interactions, and a random intercept plus random Outcome slope for each subject. To account for the non-independence between the two Outcome measurements (r = −0.325, *P* < 0.001), we specified a compound-symmetry correlation structure for the model residuals within each subject and allowed for different residual variances per Outcome. The most parsimonious fixed-effect model formula was Outcome*(Sleep/Wake*Age + Sequence).

To test whether context familiarization contributed to subsequent memory performance independently of sleep-related effects on encoding, we conducted an LMM analysis on behavior during the familiarization phase. Context error rate served as the dependent variable and was modeled as a function of the percentage of targeted play during familiarization, experimental condition (sleep vs. wake), their interaction, controlling for the transition probability between the critical behavioral states at encoding (0–1). Participant identity was included as a random intercept.

The selective effects of sleep and encoding behavior on CMI and DMI were also tested with an LMM implemented with *nlme* (**Table S6**). CMI and DMI were negatively correlated (ρ = −0.573, *P* < 0.001). We therefore included both outcome variables into one LMM. Fixed effects were Outcome (CMI vs. DMI), Sleep/Wake, Transition Probability (of behaviors 0-1), Age, and their interactions, and a random intercept plus random Outcome slope for each subject. To account for the non-independence between the two Outcome measurements, we again specified a compound-symmetry correlation structure for the model residuals within each subject and allowed for different residual variances per Outcome. The most parsimonious fixed-effect model formula was Outcome*(Age*Sleep/Wake+Transition Prob.*Sleep/Wake+Sequence).

The differences in trial-based transition probability between behaviors 0-1 across conditions and outcome measures were tested with a generalized linear mixed-effects model implemented with *afex*, including a Gamma distribution and log link function given its violation of the normality assumption (W = 0.955, *P* = 0.007). The model included the fixed effects for Outcome (context errors vs. correct trials vs. random errors), Sleep/Wake, their interaction, and a random intercept for each subject.

Whenever significant interactions were observed, we derived model-adjusted (“least-squares”) means (EMMs) using the *emmeans* package. EMMs were extracted and pairwise contrasts computed with Holm-Bonferroni adjustment for multiple comparisons to obtain simple effects.

### Sleep and Behavior Analyses

All spindle parameters (i.e. density, amplitude, duration) and SO parameters (i.e. density, amplitude, duration and slope) were compared across channels (C3, Cz, C4) using two one-way analyses of variance (ANOVAs) in MATLAB (R2023a) with Type-III sums of squares. To examine how spindle and SO characteristics predicted spatial memory performance, we fit two linear regression models in R with context-error rate as the dependent variable. The spindle and SO models both included Density, Amplitude, Duration, Channel, and the Channel × Amplitude interaction. A set of robust regressions was conducted with the CMI as the outcome using the *robustbase* package using the same set of fixed factors (**Table S7**).

SO-spindle co-occurrence rates were tested against chance level for each channel using paired-sample t-tests. For PETHs, we compared group-level bin counts against surrogate distributions using paired-sample t-tests at each time bin and channel. To correct for multiple comparisons, we employed cluster-based permutation testing (5,000 permutations).

All further analyses described here were conducted for channel Cz only. Phase–amplitude coupling was assessed by Rayleigh tests on each subject’s distribution of maximal spindle amplitudes across the SO phase cycle; significant Rayleigh statistics indicate nonuniform phase preferences. Finally, phase-amplitude coupling (vector length) was correlated with the CMI using Spearman’s ρ for those subjects with a significant Rayleigh test. We conducted an exploratory mediation analysis in R to investigate whether spindle density mediated the relationship between phase-amplitude coupling and CMI. First, a mediator model was fitted using linear regression to test the effect of spindle–SO coupling strength (predictor) on spindle density (mediator). Subsequently, an outcome model was estimated predicting the CMI from both spindle–SO coupling strength and spindle density. Mediation was formally assessed via a non-parametric bootstrap procedure (10,000 resamples) using the *mediation* package. Significance and confidence intervals of indirect (mediation) effects were estimated accordingly.

Time–frequency representations (6–20 Hz) of SO and baseline trials at Cz were contrasted for each participant via independent-samples t-tests to produce t-maps of differences. To calculate the within-subject contrasts, we generated 100 random stimulus–baseline trial pairs (preserving the N2:N3 ratio). The t-maps were tested against zero on a group level using cluster-based permutation tests (5,000 permutations) in the ±1.2-s window around the SO trough. Finally, mean t-values within the spindle band (12–16 Hz) at 400–600 ms post-trough determined by the PETHs analysis were correlated with the CMI using Spearman’s ρ; significance was determined via cluster-corrected bootstrap resampling (5,000 iterations).

## Data Availability

All main behavioral outcomes and preprocessed EEG datasets were deposited into the Open Science Framework (OSF) database and are available at the following URL (https://osf.io/yuwj5/?view_only=56945295d3a14f998ebaee909febe206).

## Code Availability

All custom computer code used to generate results reported in the manuscript that are central to the main claims are available on GitHub (https://github.com/LisaBastian/ContainerKids.git).

## Acknowledgements

We thank Tim Näher regarding the helpful comments on the manuscript. We also thank Juliana Schiebel, Jennifer Ernst, Nina Lutz, Marie Hess, Jonas Imhof and Lilian Kaim who supported data acquisition and data curation. This study was supported by grants from the Deutsche Forschungsgemeinschaft (DFG Bo 854/18-1) and the European Research Council to J.B. (ERC AdG 883098 Sleep Balance).

## Author Contributions (CRediT)

**L.B.:** Writing– original draft, Writing– review & editing, Visualization, Validation, Software, Methodology, Formal analysis, Data curation, Investigation, Conceptualization. **E.M.K.:** Writing– review & editing, Data curation, Project administration, Investigation Conceptualization. **L.G.:** Data curation, Investigation. **H.N.:** Project administration, Investigation, Conceptualization. **J.B.:** Writing– review & editing, Validation, Resources, Project administration, Funding acquisition, Conceptualization.

## References

1. Diekelmann, S. & Born, J. The memory function of sleep. Nat. Rev. Neurosci. 11, 114–126 (2010).

2. Girardeau, G. & Lopes-Dos-Santos, V. Brain neural patterns and the memory function of sleep. Science 374, 560–564 (2021).

3. Brodt, S., Inostroza, M., Niethard, N. & Born, J. Sleep-A brain-state serving systems memory consolidation. Neuron 111, 1050–1075 (2023).

4. Schapiro, A. C. et al. The hippocampus is necessary for the consolidation of a task that does not require the hippocampus for initial learning. Hippocampus 29, 1091–1100 (2019).

5. Sawangjit, A. et al. The hippocampus is crucial for forming non-hippocampal long-term memory during sleep. Nature 564, 109–113 (2018).

6. Dickey, C. W., et al. Widespread ripples synchronize human cortical activity during sleep, waking, and memory recall. Proc. Natl. Acad. Sci. 119, e2107797119 (2022).

7. Klinzing, J. G., Niethard, N. & Born, J. Mechanisms of systems memory consolidation during sleep. Nat. Neurosci. 22, 1598–1610 (2019).

8. Latchoumane, C.-F. V., Ngo, H.-V. V., Born, J. & Shin, H.-S. Thalamic Spindles Promote Memory Formation during Sleep through Triple Phase-Locking of Cortical, Thalamic, and Hippocampal Rhythms. Neuron 95, 424–435.e6 (2017).

9. Maingret, N., Girardeau, G., Todorova, R., Goutierre, M. & Zugaro, M. Hippocampo-cortical coupling mediates memory consolidation during sleep. Nat. Neurosci. 19, 959–964 (2016).

10. Staresina, B. P. et al. Hierarchical nesting of slow oscillations, spindles and ripples in the human hippocampus during sleep. Nat. Neurosci. 18, 1679–1686 (2015).

11. Rosanova, M. & Ulrich, D. Pattern-Specific Associative Long-Term Potentiation Induced by a Sleep Spindle-Related Spike Train. J. Neurosci. 25, 9398–9405 (2005).

12. Seibt, J. et al. Cortical dendritic activity correlates with spindle-rich oscillations during sleep in rodents. Nat. Commun. 8, 684 (2017).

13. Niethard, N., Ngo, H.-V. V., Ehrlich, I. & Born, J. Cortical circuit activity underlying sleep slow oscillations and spindles. Proc. Natl. Acad. Sci. 115, E9220–E9229 (2018).

14. Bastian, L. et al. Spindle–slow oscillation coupling correlates with memory performance and connectivity changes in a hippocampal network after sleep. Hum. Brain Mapp. 43, 3923–3943 (2022).

15. Kurdziel, L., Duclos, K. & Spencer, R. M. C. Sleep spindles in midday naps enhance learning in preschool children. Proc. Natl. Acad. Sci. 110, 17267–17272 (2013).

16. Werchan, D. M., Kim, J.-S. & Gómez, R. L. A daytime nap combined with nighttime sleep promotes learning in toddlers. J. Exp. Child Psychol. 202, 105006 (2021).

17. Friedrich, M., Mölle, M., Born, J. & Friederici, A. D. Memory for nonadjacent dependencies in the first year of life and its relation to sleep. Nat. Commun. 13, 7896 (2022).

18. Hermesch, N., Konrad, C., Barr, R., Herbert, J. S. & Seehagen, S. Sleep-dependent memory consolidation of televised content in infants. J. Sleep Res. 33, e14121 (2024).

19. Bastian, L. et al. Long-term memory formation for voices during sleep in three-month-old infants. Neurobiol. Learn. Mem. 215, 107987 (2024).

20. Desrochers, P. C., Kurdziel, L. B. F. & Spencer, R. M. C. Delayed benefit of naps on motor learning in preschool children. Exp. Brain Res. 234, 763–772 (2016).

21. Gómez, R. L., Bootzin, R. R. & Nadel, L. Naps Promote Abstraction in Language-Learning Infants. Psychol. Sci. 17, 670–674 (2006).

22. Mason, G. M. & Spencer, R. M. C. Sleep and Memory in Infancy and Childhood. Annu. Rev. Dev. Psychol. 4, 89–108 (2022).

23. Bevandić, J. et al. Episodic memory development: Bridging animal and human research. Neuron 112, 1060–1080 (2024).

24. Keresztes, A., Ngo, C. T., Lindenberger, U., Werkle-Bergner, M. & Newcombe, N. S. Hippocampal Maturation Drives Memory from Generalization to Specificity. Trends Cogn. Sci. 22, 676–686 (2018).

25. Mullally, S. L. & Maguire, E. A. Learning to remember: the early ontogeny of episodic memory. Dev. Cogn. Neurosci. 9, 12–29 (2014).

26. Sorrells, S. F. et al. Human hippocampal neurogenesis drops sharply in children to undetectable levels in adults. Nature 555, 377–381 (2018).

27. Nickel, K. et al. Manual morphometry of hippocampus and amygdala in adults with attention-deficit hyperactivity disorder. Psychiatry Res. Neuroimaging 267, 32–35 (2017).

28. Seress, L. & Mrzljak, L. Postnatal development of mossy cells in the human dentate gyrus: A light microscopic Golgi study. Hippocampus 2, 127–141 (1992).

29. Seress, L., Ábrahám, H., Tornóczky, T. & Kosztolányi, G. Cell formation in the human hippocampal formation from mid-gestation to the late postnatal period. Neuroscience 105, 831–843 (2001).

30. Keresztes, A. et al. Hippocampal maturity promotes memory distinctiveness in childhood and adolescence. Proc. Natl. Acad. Sci. 114, 9212–9217 (2017).

31. Huber, R. & Born, J. Sleep, synaptic connectivity, and hippocampal memory during early development. Trends Cogn. Sci. 18, 141–152 (2014).

32. Gómez, R. L. & Edgin, J. O. The extended trajectory of hippocampal development: Implications for early memory development and disorder. Dev. Cogn. Neurosci. 18, 57–69 (2016).

33. Lutz, N. D., Harkotte, M. & Born, J. Sleep’s contribution to memory formation. Physiol. Rev. 106, 363–483 (2026).

34. Jabès, A., Lavenex, P. B., Amaral, D. G. & Lavenex, P. Postnatal development of the hippocampal formation: A stereological study in macaque monkeys. J. Comp. Neurol. 519, 1051–1070 (2011).

35. García-Pérez, M. A. et al. Cortico-Hippocampal Oscillations Are Associated With the Developmental Onset of Hippocampal-Dependent Memory. Front. Neurosci. 16, (2022).

36. Fechner, J. et al. Sleep-slow oscillation-spindle coupling precedes spindle-ripple coupling during development. Sleep 47, zsae061 (2024).

37. Jaramillo, V. et al. An infant sleep electroencephalographic marker of thalamocortical connectivity predicts behavioral outcome in late infancy. NeuroImage 269, 119924 (2023).

38. Kurz, E.-M., Zinke, K. & Born, J. Sleep electroencephalogram (EEG) oscillations and associated memory processing during childhood and early adolescence. Dev. Psychol. 59, 297–311 (2023).

39. Kurz, E.-M., Bastian, L., Mölle, M., Born, J. & Friedrich, M. Development of slow oscillation–spindle coupling from infancy to toddlerhood. SLEEP Adv. 5, zpae084 (2024).

40. Hahn, M. A., Heib, D., Schabus, M., Hoedlmoser, K. & Helfrich, R. F. Slow oscillation-spindle coupling predicts enhanced memory formation from childhood to adolescence. eLife 9, e53730 (2020).

41. Knickmeyer, R. C. et al. A Structural MRI Study of Human Brain Development from Birth to 2 Years. J. Neurosci. 28, 12176–12182 (2008).

42. Utsunomiya, H., Takano, K., Okazaki, M. & Mitsudome, A. Development of the Temporal Lobe in Infants and Children: Analysis by MR-Based Volumetry. (1999).

43. Eschman, B. & Ross-Sheehy, S. Visual Short-Term Memory Persists Across Multiple Fixations: An n-Back Approach to Quantifying Capacity in Infants and Adults. Psychol. Sci. 34, 370–383 (2023).

44. Mooney, L., Dadra, J., Davinson, K., Tani, N. & Ghetti, S. Memory for space and time in 2-year-olds. Cogn. Dev. 70, 101443 (2024).

45. Newcombe, N., Huttenlocher, J., Drummey, A. B. & Wiley, J. G. The development of spatial location coding: Place learning and dead reckoning in the second and third years. Cogn. Dev. 13, 185–200 (1998).

46. Newcombe, N. S., Balcomb, F., Ferrara, K., Hansen, M. & Koski, J. Two rooms, two representations? Episodic-like memory in toddlers and preschoolers. Dev. Sci. 17, 743–756 (2014).

47. Ribordy, F., Jabès, A., Banta Lavenex, P. & Lavenex, P. Development of allocentric spatial memory abilities in children from 18 months to 5 years of age. Cognit. Psychol. 66, 1–29 (2013).

48. Sluzenski, J., Newcombe, N. S. & Kovacs, S. L. Binding, relational memory, and recall of naturalistic events: A developmental perspective. J. Exp. Psychol. Learn. Mem. Cogn. 32, 89–100 (2006).

49. Bevandić, J. et al. Episodic memory development: Bridging animal and human research. Neuron 112, 1060–1080 (2024).

50. DeMaster, D., Pathman, T. & Ghetti, S. Development of memory for spatial context: hippocampal and cortical contributions. Neuropsychologia 51, 2415–2426 (2013).

51. Lutz, N. D., Harkotte, M. & Born, J. Sleep’s contribution to memory formation. Physiol. Rev. ACCEPTED doi:ACCEPTED.

52. Noack, H., Doeller, C. F. & Born, J. Sleep strengthens integration of spatial memory systems. Learn. Mem. Cold Spring Harb. N 28, 162–170 (2021).

53. Lehnung, M. et al. The role of locomotion in the acquisition and transfer of spatial knowledge in children. Scand. J. Psychol. 44, 79–86 (2003).

54. DeLoache, J. S. & Todd, C. M. Young children’s use of spatial categorization as a mnemonic strategy. J. Exp. Child Psychol. 46, 1–20 (1988).

55. Contreras, M. P. et al. Context memory formed in medial prefrontal cortex during infancy enhances learning in adulthood. Nat. Commun. 15, 2475 (2024).

56. Hoedlmoser, K. Co-evolution of sleep spindles, learning and memory in children. Curr. Opin. Behav. Sci. 33, 138–143 (2020).

57. Klinzing, J. G. et al. Spindle activity phase-locked to sleep slow oscillations. NeuroImage 134, 607–616 (2016).

58. Helfrich, R. F., Mander, B. A., Jagust, W. J., Knight, R. T. & Walker, M. P. Old Brains Come Uncoupled in Sleep: Slow Wave-Spindle Synchrony, Brain Atrophy, and Forgetting. Neuron 97, 221–230.e4 (2018).

59. Schreiner, T., Petzka, M., Staudigl, T. & Staresina, B. P. Endogenous memory reactivation during sleep in humans is clocked by slow oscillation-spindle complexes. Nat. Commun. 12, 3112 (2021).

60. Yates, T. S. et al. Hippocampal encoding of memories in human infants. Science 387, 1316–1320 (2025).

61. Lee, J. K., Johnson, E. G. & Ghetti, S. Hippocampal Development: Structure, Function and Implications. in The Hippocampus from Cells to Systems: Structure, Connectivity, and Functional Contributions to Memory and Flexible Cognition (eds Hannula, D. E. & Duff, M. C.) 141–166 (Springer International Publishing, Cham, 2017). doi:10.1007/978-3-319-50406-3_6.

62. Ramsaran, A. I., Sanders, H. R. & Stanton, M. E. Determinants of object-in-context and object-place-context recognition in the developing rat. Dev. Psychobiol. 58, 883–895 (2016).

63. Sanders, H. R., Heroux, N. A. & Stanton, M. E. Infant rats can acquire, but not retain contextual associations in object-in-context and contextual fear conditioning paradigms. Dev. Psychobiol. 62, 1158–1164 (2020).

64. Souabni, M. et al. Napping and memory consolidation in early childhood: A systematic review and meta-analysis. Sleep Med. 133, 106649 (2025).

65. Seehagen, S., Zmyj, N. & Herbert, J. S. Remembering in the Context of Internal States: The Role of Sleep for Infant Memory. Child Dev. Perspect. 13, 110–115 (2019).

66. Joechner, A.-K., Hahn, M. A., Gruber, G., Hoedlmoser, K. & Werkle-Bergner, M. Sleep spindle maturity promotes slow oscillation-spindle coupling across child and adolescent development. eLife 12, e83565 (2023).

67. Purcell, S. M. et al. Characterizing sleep spindles in 11,630 individuals from the National Sleep Research Resource. Nat. Commun. 8, 15930 (2017).

68. Shinomiya, S., Nagata, K., Takahashi, K. & Masumura, T. Development of Sleep Spindles in Young Children and Adolescents. Clin. Electroencephalogr. 30, 39–43 (1999).

69. Kwon, H. et al. Sleep spindles in the healthy brain from birth through 18 years. Sleep 46, zsad017 (2023).

70. Seibt, J. & Frank, M. G. Primed to Sleep: The Dynamics of Synaptic Plasticity Across Brain States. Front. Syst. Neurosci. 13, 2 (2019).

71. Timofeev, I. et al. Short- and medium-term plasticity associated with augmenting responses in cortical slabs and spindles in intact cortex of cats in vivo. J. Physiol. 542, 583–598 (2002).

72. Liu, Y., Dolan, R. J., Kurth-Nelson, Z. & Behrens, T. E. J. Human Replay Spontaneously Reorganizes Experience. Cell 178, 640–652.e14 (2019).

73. Prabhakar, J., Johnson, E. G., Nordahl, C. W. & Ghetti, S. Memory-related hippocampal activation in the sleeping toddler. Proc. Natl. Acad. Sci. 115, 6500–6505 (2018).

74. Lewis, P. A. & Durrant, S. J. Overlapping memory replay during sleep builds cognitive schemata. Trends Cogn. Sci. 15, 343–351 (2011).

75. Lee, J. K., Ekstrom, A. D. & Ghetti, S. Volume of hippocampal subfields and episodic memory in childhood and adolescence. NeuroImage 94, 162–171 (2014).

76. Tamnes, C. K., Bos, M. G. N., van de Kamp, F. C., Peters, S. & Crone, E. A. Longitudinal development of hippocampal subregions from childhood to adulthood. Dev. Cogn. Neurosci. 30, 212–222 (2018).

77. Callow, D. D., Canada, K. L. & Riggins, T. Microstructural Integrity of the Hippocampus During Childhood: Relations With Age and Source Memory. Front. Psychol. 11, (2020).

78. Morris, R. G. M., Garrud, P., Rawlins, J. N. P. & O’Keefe, J. Place navigation impaired in rats with hippocampal lesions. Nature 297, 681–683 (1982).

79. Yu, J. Y. & Frank, L. M. Prefrontal cortical activity predicts the occurrence of nonlocal hippocampal representations during spatial navigation. PLOS Biol. 19, e3001393 (2021).

80. Rudy, J. W. Context representations, context functions, and the parahippocampal-hippocampal system. Learn. Mem. Cold Spring Harb. N 16, 573–585 (2009).

81. Stanton, M. E., Murawski, N. J., Jablonski, S. A., Robinson-Drummer, P. A. & Heroux, N. A. Mechanisms of context conditioning in the developing rat. Neurobiol. Learn. Mem. 179, 107388 (2021).

82. Heroux, N. A., Horgan, C. J., Pinizzotto, C. C., Rosen, J. B. & Stanton, M. E. Medial prefrontal and ventral hippocampal contributions to incidental context learning and memory in adolescent rats. Neurobiol. Learn. Mem. 166, 107091 (2019).

83. Gómez, R. L., Bootzin, R. R. & Nadel, L. Naps Promote Abstraction in Language-Learning Infants. Psychol. Sci. 17, 670–674 (2006).

84. Hupbach, A., Gomez, R. L., Bootzin, R. R. & Nadel, L. Nap-dependent learning in infants. Dev. Sci. 12, 1007–1012 (2009).

85. Friedrich, M., Wilhelm, I., Born, J. & Friederici, A. D. Generalization of word meanings during infant sleep. Nat. Commun. 6, 6004 (2015).

86. Bowman, C. R. & Zeithamova, D. Abstract Memory Representations in the Ventromedial Prefrontal Cortex and Hippocampus Support Concept Generalization. J. Neurosci. Off. J. Soc. Neurosci. 38, 2605–2614 (2018).

87. Samborska, V., Butler, J. L., Walton, M. E., Behrens, T. E. J. & Akam, T. Complementary task representations in hippocampus and prefrontal cortex for generalizing the structure of problems. Nat. Neurosci. 25, 1314–1326 (2022).

88. Antonenko, D., Diekelmann, S., Olsen, C., Born, J. & Mölle, M. Napping to renew learning capacity: enhanced encoding after stimulation of sleep slow oscillations. Eur. J. Neurosci. 37, 1142–1151 (2013).

89. Németh, D. et al. Optimizing the methodology of human sleep and memory research. Nat. Rev. Psychol. 3, 123–137 (2024).

90. Randler, C., Faßl, C. & Kalb, N. From Lark to Owl: developmental changes in morningness-eveningness from new-borns to early adulthood. Sci. Rep. 7, 45874 (2017).

91. Tamis-LeMonda, C. S. & Masek, L. R. Embodied and Embedded Learning: Child, Caregiver, and Context. Curr. Dir. Psychol. Sci. 32, 369–378 (2023).

92. Berhenke, A., Miller, A. L., Brown, E., Seifer, R. & Dickstein, S. Observed emotional and behavioral indicators of motivation predict school readiness in Head Start graduates. Early Child. Res. Q. 26, 430–441 (2011).

93. Weber, F. D. et al. Coupling of gamma band activity to sleep spindle oscillations – a combined EEG/MEG study. NeuroImage 224, 117452 (2021).

94. Donoghue, T. et al. Parameterizing neural power spectra into periodic and aperiodic components. Nat. Neurosci. 23, 1655–1665 (2020).

95. Moca, V. V., Bârzan, H., Nagy-Dăbâcan, A. & Mureșan, R. C. Time-frequency super-resolution with superlets. Nat. Commun. 12, 337 (2021).

