## Supplementary Information for "Sleep forms flexible context representations in toddlers"

This PDF file includes:
Supplementary Methods
Supplementary Notes
Supplementary Figure 1 to 4
Supplementary Table 1 to 7

### Supplementary Methods

#### Graph neural network (GNN) analysis

We used a first-order Markov chain to model transition probabilities between all target behaviors (**Table S1**). Each behavior represents a state. Transitions were evaluated at each sample *t,* as the probability that the system moves from its current state i to a next state $j$ with

$$P_{ij} = P(X_{t+1} = j | X_t = i).$$

Collectively, these probabilities form a transition probability matrix. The rows sum to 1 as any behavior transitions to another. Each entry $(i, j)$ is the estimated probability of going from behavior $i$ to behavior $j$ in one sample $t$. Given a sequence of behaviors over time

$$X_1, X_1, \dots, X_T,$$

we counted how often each ordered pair $(i \rightarrow j)$ occurs and normalized by the total number of transitions initiated by state $i$:

$$P_{ij} = \frac{\text{count of transitions } i \rightarrow j}{\sum_k \text{count of transitions } i \rightarrow k}$$

The transition probability matrices and associated metadata were converted into graph objects. Specifically, we constructed directed graphs for each participant during familiarization of the two rooms and in each memory task trial during encoding. Target behaviors served as nodes and the directed edges were the transition probabilities between ordered pairs of behaviors. Metadata were processed to define node features (i.e. actors corresponding to each behavior, whether the behavior was compliant vs. non-compliant with the task, duration of each behavior in %), and global graph-level features (i.e. condition, session, location, room, sex, scorer, trial duration, time to target container, motivation, age and MDI). Categorical variables were encoded using one-hot representations, and continuous variables were normalized.

We, then, built a GNN to classify sleep versus wake conditions based on the overall behavior during the familiarization and encoding of the memory task. The GNN architecture utilized edge-conditioned convolutions (NNConv) to effectively integrate node and edge information: (1) The initial convolutional layer transformed input node features into embeddings of a specified dimension $D$ (8, for optimization see **Fig. 3b**). Edge attributes were mapped via a multi-layer perceptron (MLP) to convolutional weight matrices. This was followed by two additional edge-conditioned layers maintaining embedding dimensionality $D$. Activation functions were ReLU6 throughout. (2) Graph embeddings were generated by concatenating global max-pooling and mean-pooling of node embeddings, combined with global graph-level features. A fully-connected classification head with one hidden layer (128 neurons, ReLU activation) output binary class predictions through a logistic output neuron.

For GNN training and evaluation, the graphs were each split using stratified sampling, allocating 90% of graphs for model training and validation, and 10% reserved for final testing. The 90% subset was further subdivided for 10-fold cross-validation using stratified splits to preserve class distributions. Optimization utilized the Adam algorithm (learning rate $1 \times 10^{-4}$, weight decay $0.01$), supervised by binary cross-entropy loss (BCEWithLogitsLoss). Training employed gradient clipping (norm ≤ 1.0) and a learning rate scheduler (ReduceLROnPlateau) monitoring validation loss. Early stopping criteria (patience of 100 epochs without improvement) limited training to a maximum of 4000 epochs. Post-validation, models were retrained on the entire training subset for 3000 epochs and evaluated on the previously held-out 10% testing subset to report final accuracy, specificity and sensitivity of the model. Model performance was tested against an empirical chance level creating a surrogate distribution by shuffling the class labels 5000 times.

To interpret the graph neural network predictions, we applied the GNNExplainer algorithm via PyTorch Geometric's Explainer API: Node-level explanations were computed by generating masks that identify influential edges. The top 20% most significant nodes and their connections per graph were selected as the most important subgraphs for the predictions of the GNN.

**Control analyses**

To ensure that any observed differences between sleep and wake conditions could not be attributed to non-specific factors, we conducted a series of control analyses on task engagement and motivation measures. All analyses were performed in R (Version 2023.12.1) using an (Bonferroni corrected) alpha level of 0.05; effect sizes of all t-tests were reported as Cohen's d.

As an indicator of the child's motivation and sleepiness we computed the number of trials each subject completed in the sleep versus wake conditions using two-tailed paired-samples t-tests. To rule out differential exposure to the contexts during the encoding (hide) phase of the hide-and-seek game, we compared the total time (in s) each toddler spent exploring the contexts during the encoding phase across conditions. Search efficiency at retrieval was assessed by measuring the latency (in seconds) from entering the room to opening the target container. Number of trials, time at encoding, time until target and motivation ratings obtained from the video recordings were compared between sleep and wake conditions with two-tailed paired sample t-tests. To verify that performance was not influenced by sex, we conducted independent-samples t-tests comparing girls and boys on percentages of correct trials, context errors, and random errors.

An influence of increasing sleepiness on performance over time was analyzed by comparing the motivation ratings across trials using two-way ANOVAs (Sleep/Wake x Trial) for both, encoding and retrieval phases of the hide-and-seek task. To control for time-of-day effects, an independent samples t-test was performed to compare context error rates for wake participants tested in the morning (N = 15) versus afternoon (N = 13). To assess differences in retention interval between conditions we performed paired samples t-tests (N = 26). Finally, exploratory Spearman correlations (uncorrected for multiple comparisons) were calculated on the association between the time in different sleep stages and percent context errors and CMI values.

**Spindle detection**

For spindle detection, the EEG from C3, Cz, and C4 was then band-pass filtered around this personalized peak (±2 Hz, 4th-order zero-phase Butterworth). The root-mean-square (RMS) envelope of the filtered signal was computed at each sample and smoothed with a 200 ms moving average; candidate spindles were those 0.5–3 s epochs in which the smoothed RMS exceeded its mean by 1.5 SD. Peaks and troughs in each spindle were marked by the local extrema of the filtered signal, with the largest trough defining the spindle's central time point and the trough-to-peak voltage difference taken as its amplitude. Individual spindle frequency was calculated as the total number of half-cycles (peaks + troughs) divided by twice the spindle duration; adjacent spindles separated by less than 0.25 s were merged.

**Slow Oscillation (SO) detection**

SOs were detected at using a zero-phase, 6th-order Butterworth low-pass filter (cutoff 4 Hz). The continuous NonREM signal was segmented into half-waves by zero crossings, and candidate SOs were defined as negative half-waves followed by positive half-waves with oscillation frequencies of 0.3–1.25 Hz. To ensure robust detection, each SO's amplitude and trough depth had to exceed the channel's mean SO amplitude or down-peak potential, respectively. SO amplitude and frequency were then quantified using the same half-wave counting procedure as for spindles.

### Supplementary Notes

#### Supplementary Note 1. Control analyses on sex, motivation, context exposure, time-of-day effects and break duration.

Control analyses for unspecific effects of our manipulation showed that sleep and wake conditions did not differ in the number of completed trials ($t_{27}$ = -0.691, *P* = 0.500, *d* = 0.133, 95% CI [-0.67 – 0.40]), time spent in a context during encoding of the task ($t_{27}$ = 0.112, *P* = 0.912, *d* = 0.034, 95% CI [-0.59 – 0.49]), time to reach the target container at retrieval ($t_{26}$ = 1.215, *P* = 0.235, *d* = 0.298, 95% CI [-0.25 – 0.85]), or rated motivation by the experimenter during encoding ($t_{26}$ = -0.822, *P* = 0.418, *d* = -0.158, 95% CI [-0.71 – 0.39]) and retrieval ($t_{27}$ = -0.602, P = 0.539, *d* = -0.189, 95% CI [-0.74 – 0.36], see **Fig. S1**, for a detailed analysis of motivation ratings across trials). Female and male participants appeared to score similarly on the three outcome measures (percent correct: $t_{54}$ = -0.002, *P* = 0.999, *d* < 0.001, 95% CI [-0.54 – 0.54]; context error rate: $t_{54}$ = -0.693, *P* = 0.491, *d = -0.187*, 95% CI [-0.73 – 0.36]; random error rate: $t_{54}$ = 0.556, *P* = 0.580, *d* = 0.150, 95% CI [-0.39 – 0.69]).

To assess whether differences in prior wake duration or time-of-day contributed to the observed memory effects, we performed additional control analyses. Prior wake duration before familiarization was entered as a covariate into the linear mixed-effects models for context error rate, CMI, and DMI. Prior wake duration neither showed a main effect on context error rate (β = 0.06, SE = 0.06, *P* = 0.315, 95% CI [-0.07 0.14]), or CMI/DMI (β < -0.001, SE = 0.001, *P* = 0.305, 95% CI [-0.002 0.001]), nor did it interact with the Sleep/Wake condition in any model (all Sleep/Wake x Wake Duration interaction *P* > 0.435). In addition, memory performance within the wake condition did not differ between children tested in the morning (n = 15) versus the afternoon (n = 13). Independent-samples comparisons revealed no significant differences for context error rate ($t_{22}$ = 0.31, *P* = 0.765, 95% CI [-10.89 14.61], d = 0.13). Together, these analyses suggest that the observed sleep-related differences in contextual memory cannot be explained by prior wake duration or time-of-day effects.

We performed additional control analyses to examine whether differences in retention interval duration contributed to the observed memory effects. The total break duration was numerically longer in the sleep (159.42 min) than in the wake condition (134.63 min). However, a direct comparison of the effective retention interval between conditions (excluding two participants with missing timing data) revealed only a statistical trend that did not reach significance ($t_{25}$ = 1.99, *P* = 0.058; 95% CI [-0.9, 53.21]). We furthermore included etention interval duration as a covariate in the linear mixed-effects models predicting memory performance. Retention interval duration did not predict the context error rate (β = −0.03, SE = 0.05, *P* = 0.520, 95% CI [-0.13, 0.07]). Importantly, the effect of Sleep/Wake condition on context error rate remained significant after controlling for retention interval duration (β = 103.98, SE = 45.76, *P* = 0.025, 95% CI [22.99, 207.39]). Similarly, retention interval duration did not significantly predict the CMI/DMI (β = 0.001, SE = 0.001, *P* = 0.113, 95% CI [-0.001, 0.003]). Together, these analyses indicate that variation in retention interval duration does not account for the observed sleep-related effects on contextual memory.

### Supplementary Figures

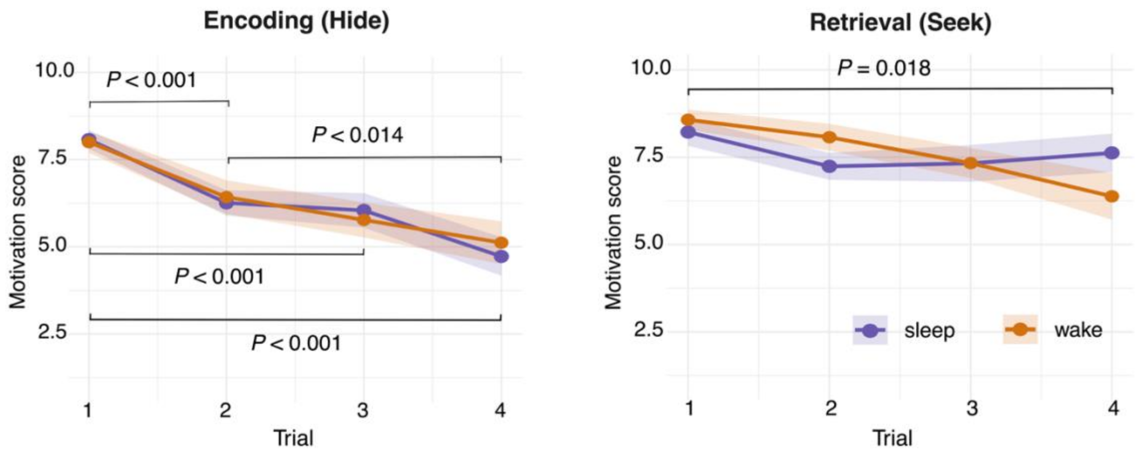


**Figure S1. Motivation scores separately for each of the 4 retrieval trials of the hide-and- seek spatial memory task.** Mean (±SEM) motivation scores for the sleep (purple) and wake (orange) condition, separately at encoding (left) and retrieval (right). *P*-values indicate significance for the follow-up t-tests (Bonferroni-corrected. Motivation scores decreased across trials in both the sleep and wake conditions during encoding ($F_{184,3}$ = 17.182, *P* < 0.001) and retrieval ($F_{184,3}$ = 3.439, *P* = 0.018) with the decreases being highly comparable between sleep and wake conditions (all *P* > 0.717) or in the interaction with trials (all P > 0.153).

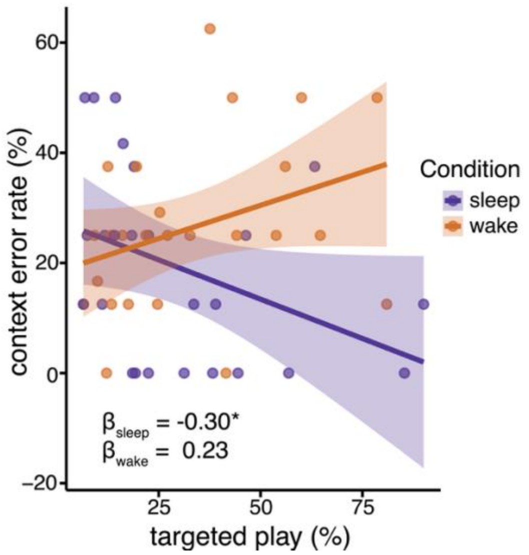


**Figure S2. Distinct effects of context familiarization on context error rate after sleep and wakefulness.** Mean (±SEM) for the effect of targeted play during context familiarization (i.e. playing with context relevant items; see Table S1) on context error rate for the sleep (purple) and wake (orange) condition. Engaging in more targeted play (relative to the trial duration) during context familiarization was associated with fewer context errors after sleep ($P$ = 0.048) and no effect after wakefulness ($P$ = 0.139). Individual data points are overlayed.

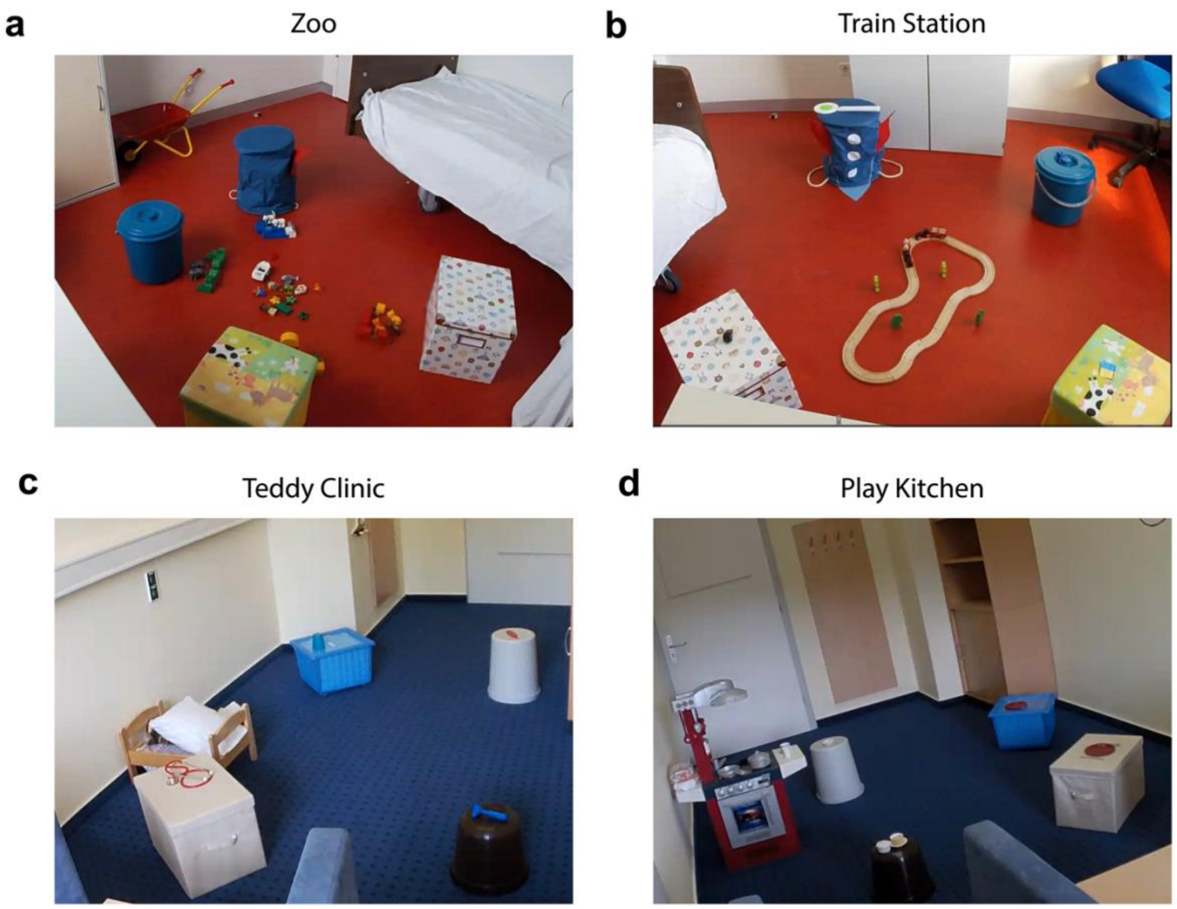


**Figure S3. Arrangement of the four spatial contexts.** Contexts **a** and **b** are in one lab (lab B) and contexts **c** and **d** in the other lab (lab A) in the city of Tübingen. The rooms were each equipped with four containers that served as hiding places for the puppets during the spatial memory task. The containers were the same for the two rooms of one condition (**a,b** or **c,d**) but arranged differently per room. In one corner of each room a chair was positioned for the parent. Each room was decorated according to a specific theme, with one big central item, another item the child could use or put on, and smaller context-specific items at each container. **a,** The zoo included a big wheelbarrow and animal enclosures for giraffes, elephants, ice bears and lions in front of the containers. A play car was positioned in the center with a zookeeper and food for the animals. **b,** The train station included a conductor's hat and wooden train tracks with a train in the center. The items on the containers were a whistle, a trowel, train tickets and a ticket validator. **c,** The teddy clinic included a teddy bear in a bed and a doctor's kit next to it. A statoscope, an otoscope, a knee hammer and a fever monitor were placed on the containers. **d,** The play kitchen was equipped with an apron and a kitchen including an oven and a stove with food and pots. Different cutlery and dishes were placed on the containers.

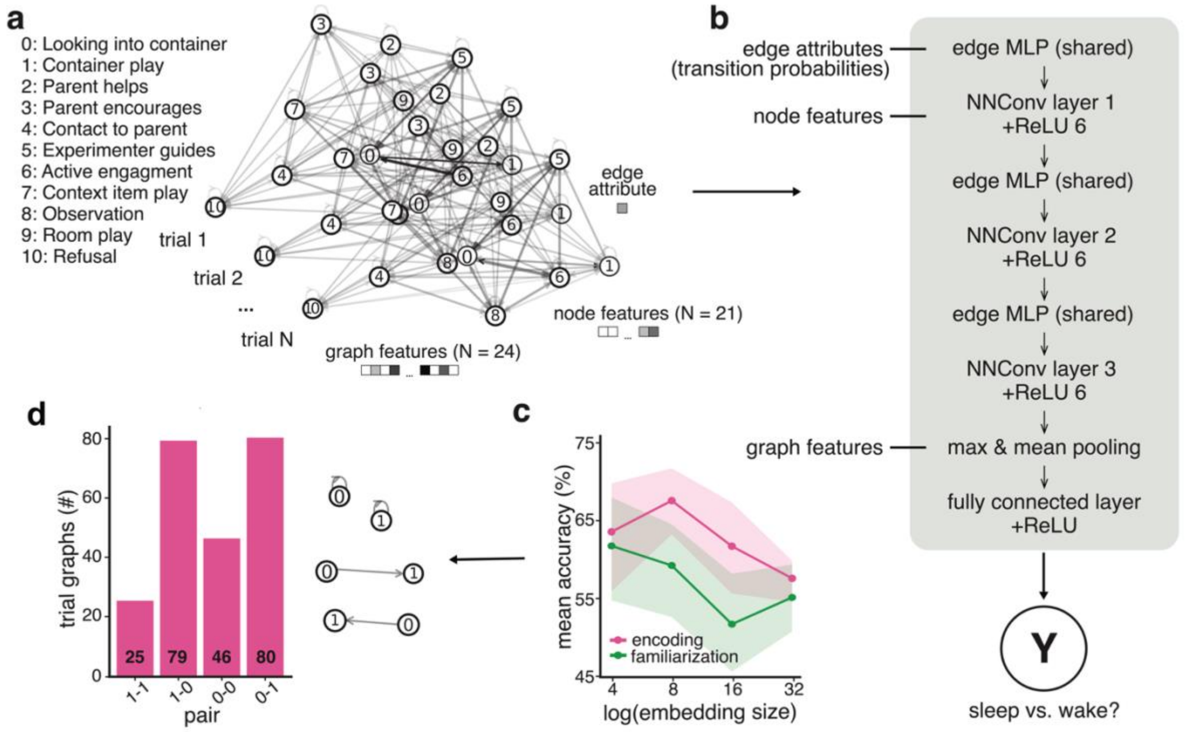


**Figure S4. Graph neural network analysis, embedding size comparison and node pair relevance.** **a,** Trial-wise directed encoding graphs constructed from video-scored behavioral sequences. Nodes represent distinct behavioral states (labels 0–10), edges indicate transition probabilities between behaviors, and node- and graph-level features encode behavioral, task-related, and demographic information. One graph was generated per trial. **b,** Graph neural network (GNN) architecture used for classification. Node features were processed by three edge-conditioned convolutional layers (NNConv) in which edge attributes (transition probabilities) were transformed into convolutional weight matrices via a shared multilayer perceptron (MLP). ReLU6 activations were applied after each convolution. Node embeddings were aggregated using global mean and max pooling and concatenated with graph-level features before classification by a fully connected layer with ReLU activation, yielding a binary prediction (sleep vs. wake). **c,** Mean (±SEM) 10-fold cross-validation (CV) accuracies across embedding sizes (i.e. the dimensionality of the latent graph representations) of the GNN at familiarization (green) and encoding (pink). The embedding size with the highest CV-accuracy (familiarization: 4, encoding: 8) was used for the final test. **d,** Histograms of the top 20% node pairs (illustrated on the right) in the graphs of each trial used for the GNN predictions at encoding. The node pair most often identified to differentiate between sleep and wake conditions was 0-1.

### Supplementary Tables

*Table S1.* Target behaviors for the video scoring analysis

| *No.* | *Event type* | *Actor* | *Task phase* | *Behavior* | *Description* |
|---|---|---|---|---|---|
| X | point | child | fam./enc. | room entrance | Child places the first foot in the room |
| X | point | child | fam./enc. | room exit | Child places the first foot out of the room |
| X | point | child | enc. | target container | Child opens the container where the puppet is to be hidden |
| 0 | state | child | fam./enc. | looking | Child looks into the container |
| 1 | state | child | fam./enc. | container play | Child is playing with a container |
| 2 | state | parent/ exp. | enc. | help | Parent/experimenter gives unallowed hints as to where the puppet is hidden |
| 3 | state | parent | fam./enc | contact parent | Parent encourages child to continue with the task |
| 4 | state | child | fam./enc. | contact child | Child seeks contact to the parent, hoping for help |
| 5 | state | exp. | fam./enc. | instructing | Experimenter indicates the behavior that is expected from the child to encourage |
| 6 | state | child | fam./enc. | engagement | Child follows the experimenter's instruction |
| 7 | state | child | fam./enc. | context play | Child is distracted from the task, playing with the objects related to the context |
| 8 | state | child | fam./enc. | observation | Child is observing while the experimenter/parent provide guidance |
| 9 | state | child | fam./enc | uncoordinated play | Child is distracted from the task, playing with context-unrelated objects |
| 10 | state | child | fam./enc. | refusal | Child refuses to do anything anymore |
| 11 | state | child | fam. | creative play | At the end the context familiarization, the child plays freely in the context |
| 12 | state | child | fam. | targeted play | Child plays with items unique to the context |

*Note.* fam. = familiarization, enc. = encoding, exp. = experimenter, Point events occur at one specific timepoint. State events have a start and endpoint.

*Table S2.* Sleep Macroarchitecture (*N* = 25)

| *Parameter* | *Mean (SD)* | *range* |
|---|---|---|
| TST (min) | 92.94 (30.74) | 27.00 – 156.50 |
| WASO (min) | 1.36 (2.98) | 0.00 – 10.00 |
| N1 (min) | 1.20 (1.24) | 0.00 – 4.00 |
| N2 (min) | 22.88 (15.65) | 3.50 – 56.00 |
| N3 (min) | 60.64 (21.68) | 22.50 – 91.50 |
| REM (min) | 8.22 (8.34) | 0.00 – 28.50 |
| WASO (% of TST) | 1.29 (2.78) | 0.00 – 11.36 |
| N1 (% of TST) | 1.42 (1.53) | 0.00 – 5.33 |
| N2 (% of TST) | 23.73 (12.59) | 4.71 – 50.50 |
| N3 (% of TST) | 65.94 (14.55) | 41.46 – 91.48 |
| REM (% of TST) | 8.90 (8.20) | 0.00 – 21.54 |

*Note.* TST = total sleep time, WASO = wake after sleep onset, REM = rapid eye movement

*Table S3.* Characteristics of spindle and SO events (*N* = 25)

| | SOs | | | Spindles | | |
|---|---|---|---|---|---|---|
| | *C3* | *Cz* | *C4* | *C3* | *Cz* | *C4* |
| Mean frequency (Hz) | --- | --- | --- | 12.97 (0.06) | 12.83 (0.05) | 12.94 (0.07) |
| Down-to-up slope (µV / s) | 139.78 (11.13) | 168.56 (13.10) | 136.77 (10.72) | --- | --- | --- |
| Density (#/min) | 4.16 (0.14) | 4.45 (0.10) | 4.27 (0.10) | 2.98 (0.12) | 3.37 (0.17) | 3.08 (0.17) |
| Amplitude (µV) | 296.10 (10.69) | 361.96 (11.22) | 304.62 (10.07) | 31.77 (1.42) | 38.78 (1.76) | 31.70 (1.43) |
| Duration (s) | 1.14 (0.01) | 1.18 (0.02) | 1.19 (0.03) | 0.81 (0.01) | 0.83 (0.01) | 0.81 (0.01) |

*Note.* Values expressed as mean (SD).

*Table S4.* Sleep Stage Correlations with Context Memory Measures (*N* = 25)

| | Context Errors | | CMI | |
|---|---|---|---|---|
| Parameter | *Pearson r* | *P-value** | *Spearman* ρ | *P-value** |
| TST | 0.119 | 0.412 | -0.160 | 0.266 |
| WASO (% of TST) | -0.072 | 0.618 | 0.153 | 0.288 |
| N1 (% of TST) | -0.246 | 0.236 | 0.214 | 0.304 |
| N2 (% of TST) | 0.182 | 0.384 | -0.191 | 0.361 |
| N3 (% of TST) | -0.185 | 0.376 | 0.123 | 0.559 |
| REM (% of TST) | 0.094 | 0.654 | -0.065 | 0.757 |

*Note.* TST = total sleep time, WASO = wake after sleep onset, REM = rapid eye movement, * uncorrected for multiple comparisons

*Table S5.* Fixed effects of the linear-mixed effects model analysis for the selective effect of sleep on context error rate vs. percent correct trials controlling for age (in days) and room sequence (N = 28)

| DV | Predictor | *β* | *SE* | *DF* | *t* | *P* |
|---|---|---|---|---|---|---|
| memory outcome value (%) | intercept | 39.70 | 45.22 | 186 | 0.88 | 0.381 |
| | outcome (context error rate vs. %-correct trials) | -62.18 | 56.03 | 186 | -1.11 | 0.269 |
| | condition (sleep vs. wake) | -100.10 | 63.52 | 186 | -1.58 | 0.117 |
| | age (in months) | -0.49 | 1.50 | 26 | -0.33 | 0.745 |
| | **sequence (same vs. different)** | **14.34** | **5.48** | **186** | **2.62** | **0.009** |
| | condition * age | 3.30 | 2.15 | 186 | 1.54 | 0.126 |
| | **outcome*condition** | **205.30** | **78.00** | **186** | **2.63** | **0.009** |
| | outcome*age | 2.47 | 1.86 | 186 | 1.32 | 0.187 |
| | **outcome*sequence** | **-25.90** | **6.73** | **186** | **-3.85** | **<0.001** |
| | **outcome*condition* age** | **-6.59** | **2.64** | **186** | **-2.50** | **0.013** |

*Note.* DV = dependent variable

*Table S6***.** Fixed effects of the linear-mixed effects model analysis for the selective effect of sleep and transition probabilities (0-1) on CMI vs. DMI controlling for age (in days) and room sequence (N = 27)

| DV | Predictor | *β* | *SE* | *DF* | *t* | *P* |
|---|---|---|---|---|---|---|
| memory index value | **intercept** | 0.98 | 0.54 | 161 | 1.81 | 0.072 |
| | outcome (DMI vs. CMI) | -1.09 | 0.93 | 161 | -1.17 | 0.243 |
| | **condition (sleep vs. wake)** | < -0.001 | 0.001 | 26 | -0.90 | 0.377 |
| | age (in months) | **-1.86** | **0.75** | **161** | **-2.50** | **0.014** |
| | transition prob. | -0.40 | 0.53 | 161 | -0.75 | 0.453 |
| | **sequence (same vs. different)** | **0.17** | **0.06** | **161** | **2.83** | **0.005** |
| | outcome * age | 0.001 | 0.001 | 161 | 1.24 | 0.217 |
| | **outcome*condition** | **3.48** | **1.16** | **161** | **2.99** | **0.003** |
| | **condition*age** | **0.001** | **< 0.001** | **161** | **2.15** | **0.033** |
| | condition*transition prob. | 1.48 | 0.78 | 161 | 1.91 | 0.057 |
| | outcome*transition prob. | 0.51 | 0.92 | 161 | 0.55 | 0.580 |
| | **outcome*sequence** | **-0.30** | **0.10** | **161** | **-3.04** | **0.003** |
| | **outcome*condition* age** | **-0.003** | **0.001** | **161** | **-2.46** | **0.015** |
| | **outcome*condition* transition prob.** | **-3.70** | **1.33** | **161** | **-2.77** | **0.006** |

*Note.* DV = dependent variable

*Table S7.* Robust regression for the association between spindle/SO parameters and CMI (N = 25)

| Event | DV | Predictor | *β* | *SE* | *t* | *P* |
|---|---|---|---|---|---|---|
| Spindles | CMI | Intercept | 0.932 | 0.584 | 1.60 | 0.116 |
| | | **density (#/min)** | **0.114** | **0.044** | **2.60** | **0.012** |
| | | duration (s) | -0.211 | 0.798 | -0.27 | 0.792 |
| | | channel (C4 vs. C3) | -0.066 | 0.318 | -0.21 | 0.835 |
| | | channel (Cz vs. C3) | 0.105 | 0.301 | 0.35 | 0.728 |
| | | max. amplitude (µV) | -0.011 | 0.007 | -1.57 | 0.121 |
| | | cax. amplitude × channel (C4) | 0.002 | 0.009 | 0.26 | 0.799 |
| | | Max. amplitude × channel (Cz) | -0.001 | 0.008 | -0.15 | 0.884 |
| Slow oscillaions | CMI | Intercept | 0.184 | 1.856 | 0.10 | 0.922 |
| | | channel (C4 vs. C3) | 2.031 | 2.873 | 0.71 | 0.483 |
| | | channel (Cz vs. C3) | -2.046 | 2.209 | -0.93 | 0.359 |
| | | density (#/min) | 0.037 | 0.116 | 0.32 | 0.751 |
| | | duration (s) | 0.396 | 1.435 | 0.28 | 0.784 |
| | | amplitude (µV) | 0.002 | 0.006 | 0.38 | 0.709 |
| | | slope (µV/s) | 0.00054 | 0.00121 | 0.44 | 0.661 |
| | | density × channel (C4) | -0.067 | 0.180 | -0.37 | 0.710 |
| | | density × channel (Cz) | 0.226 | 0.152 | 1.49 | 0.142 |
| | | duration × channel (C4) | -1.671 | 2.026 | -0.83 | 0.413 |
| | | duration × channel (Cz) | 1.078 | 1.626 | 0.66 | 0.510 |
| | | amplitude × channel (C4) | 0.00347 | 0.00756 | 0.46 | 0.648 |
| | | amplitude × channel (Cz) | -0.00784 | 0.00828 | -0.95 | 0.348 |
| | | slope × channel (C4) | 0.00062 | 0.00157 | 0.39 | 0.696 |
| | | slope × channel (Cz) | -0.00153 | 0.00163 | -0.94 | 0.352 |

*Note.* CMI = contextualized memory index, DV = dependent variable, C3 served as reference channel
